# Rapid and transient evolution of local adaptation to seasonal host fruits in an invasive pest fly

**DOI:** 10.1101/2022.03.01.482503

**Authors:** Laure Olazcuaga, Julien Foucaud, Candice Deschamps, Anne Loiseau, Jean-Loup Claret, Romain Vedovato, Robin Guilhot, Cyril Sevely, Mathieu Gautier, Ruth A. Hufbauer, Nicolas O. Rode, Arnaud Estoup

## Abstract

Both local adaptation and adaptive phenotypic plasticity can influence the match between phenotypic traits and local environmental conditions. Theory predicts that environments stable for multiple generations promote local adaptation, while highly heterogeneous environments favor adaptive phenotypic plasticity. However, when environments have periods of stability mixed with heterogeneity, the relative importance of local adaptation and adaptive phenotypic plasticity is unclear. Here, we used *Drosophila suzukii* as a model system to evaluate the relative influence of genetic and plastic effects on the match of populations to environments with periods of stability from three to four generations. This invasive pest insect can develop within different fruits, and persists throughout the year in a given location on a succession of distinct host fruits, each one being available for only a few generations. Using reciprocal common environment experiments of natural *D. suzukii* populations collected from cherry, strawberry and blackberry, we found that both oviposition preference and offspring performance were higher on medium made with the fruit from which the population originated, than on media made with alternative fruits. This pattern, which remained after two generations in the laboratory, was analyzed using a statistical method we developed to quantify the contributions of local adaptation and adaptive plasticity in determining fitness. Altogether, we found that genetic effects (local adaptation) dominate over plastic effects (adaptive phenotypic plasticity). Our study demonstrates that spatially and temporally variable selection does not prevent the rapid evolution of local adaptation in natural populations. The speed and strength of adaptation may be facilitated by several mechanisms including a large effective population size and strong selective pressures imposed by host plants.

**Impact Summary:** Natural populations often exhibit good “fit” to the environment they are in. However, environments change over time and space, and following change, the fit between a population and its environment may be poor. A question of long-standing interest is how do populations track changing environments to maintain fitness? Two main mechanisms are known: (*i*) genetic shifts in the form of local adaptation, in which traits evolve over time through differences in fitness of individuals harboring different genetic variants, and (*ii*) plastic shifts, or adaptive phenotypic plasticity, in which traits immediately change in response to environmental change. Adaptation is common when environments change over multiple generations, while plasticity is common when environments change over an individual’s lifetime. However, it remains unclear whether plasticity or adaptation is more vital to maintaining fitness when environments change at an intermediate pace.

*Drosophila suzukii* is well-suited to evaluating the relative importance of plasticity and adaptation in response to an intermediate pace of environmental change. This invasive pest species experiences an environment that shifts every 1-4 generations as host fruits change over time and space. Here, we studied natural populations of *D. suzukii* collected from different hosts. Using reciprocal common environment experiments, we evaluated their fitness on their source and alternative hosts.

*Drosophila suzukii* populations were most fit on their source host, successfully tracking an intermediate pace of environmental change. We developed a statistical method to quantify the contributions of adaptive plasticity and local adaptation in determining fitness. We found that fitness was mainly maintained through local adaptation to each new host in succession. This study highlights that spatially and temporally variable selection does not prevent local adaptation and, on the contrary, illustrates how rapid the adaptive process can be. It also provides a novel statistical tool that can be applied to other systems

## Introduction

Adaptation is the process whereby organisms come to match environmental conditions in ways that enhance fitness. When selection differs between environments, adaptation to the home environment may result in reduced fitness in other environments (e.g., Via, 1991; Torres Dowdall *et al*., 2012, Kawecki & Ebert, 2004). Such a pattern of higher fitness in the natal environment than in alternate environments is typically understood to be due to local adaptation of genetically differentiated populations. However, this pattern can also arise through adaptive phenotypic plasticity (Torres Dowdall *et al*., 2012; Yampolsky *et al*., 2014; Rago *et al*., 2019; Bonnet *et al*., 2021; Enbody *et al*., 2021), which enables populations to rapidly track environmental change without genetic change, through shifts in behavior and development, or parental and trans-generational effects (Price *et al*., 2003).

Adaptive phenotypic plasticity can modify phenotypes more rapidly than adaptation from standing genetic variation (Levins, 1968; Gillespie, 1974; Botero *et al*., 2015; Tufto, 2015). It is expected to evolve when environmental change is frequent, occurring within a generation, and can be assessed through reliable cues such as changing light regimes predicting oncoming colds (Gavrilets & Scheiner, 1993; Jong, 1999; Tufto, 2000). However, plasticity may be costly, in that it requires energy to alter phenotypes, in addition to the material expenses involved in sensory and regulatory machinery (Dewitt *et al*., 1998; Van Buskirk & Steiner, 2009; Auld *et al*., 2010). Given such potential costs, when environments change more slowly, i.e., over the course of multiple generations, genetic differentiation leading to local adaptation is favored by selection (Levins, 1968).

Many environments change at an intermediate frequency, and are not necessarily predictable. Population responses to such change could be shaped by genetic adaptation, phenotypic plasticity, or both (Schmid, 1992). Here, we seek to understand the relative contribution of local adaptation and adaptive plasticity in maintaining fitness of populations living in anthropogenically altered environments, specifically in agricultural areas, that change at an intermediate frequency.

Local adaptation can be distinguished from adaptive phenotypic plasticity by performing reciprocal common environment experiments, in which the performance of populations in their original environment as well as in other environments is measured over several generations (Fig. 1A *vs*. 1B). The relative importance of genetic and plastic responses in matching the phenotype of natural populations to their environment is just beginning to be studied in cases with frequent or infrequent environmental changes (Rago *et al*., 2019; Bonnet *et al*., 2021; Enbody *et al*., 2021), or in seasonal changes (Stone *et al*., 2019). However, to our knowledge, no study has investigated the importance of local adaptation and phenotypic plasticity when populations evolve in heterogeneous environments with periods of environmental stability from three to four generations. Three main experimental and statistical reasons account for an overall lack of empirical evidence regarding the relative importance of genetic and plastic responses in this situation. First, selective pressures are hard to control *in natura*, as they can vary both spatially and temporally (Rausher, 1988; Fry, 1996; Hansen *et al*., 2006; Barghi *et al*., 2020). This issue is best tackled by performing reciprocal common environment experiments in the laboratory or other controlled environment where local adaptation to a given biotic or abiotic factor of interest can be tested on populations from different geographical locations (Fig. 1A; Turesson, 1922; Kawecki & Ebert, 2004; Hereford, 2009). Second, local adaptation can be masked by variation in performance among populations due to environmental effects (non-adaptive phenotypic plasticity; Fig. 1C). Indeed, local adaptation can only be distinguished from non-adaptative and adaptive phenotypic plasticity by performing reciprocal common environment experiments. In these experiments, individuals from populations from different geographical locations are raised in the same environment over one or several generations (Kawecki & Ebert, 2004). Third, although multi-generation laboratory experiments allow estimates of both genetic and plastic responses (Merila & Hendry, 2014), reliable statistical tools to quantify and test their relative contribution to the performance of populations across environments are currently lacking.

**Figure 1.**
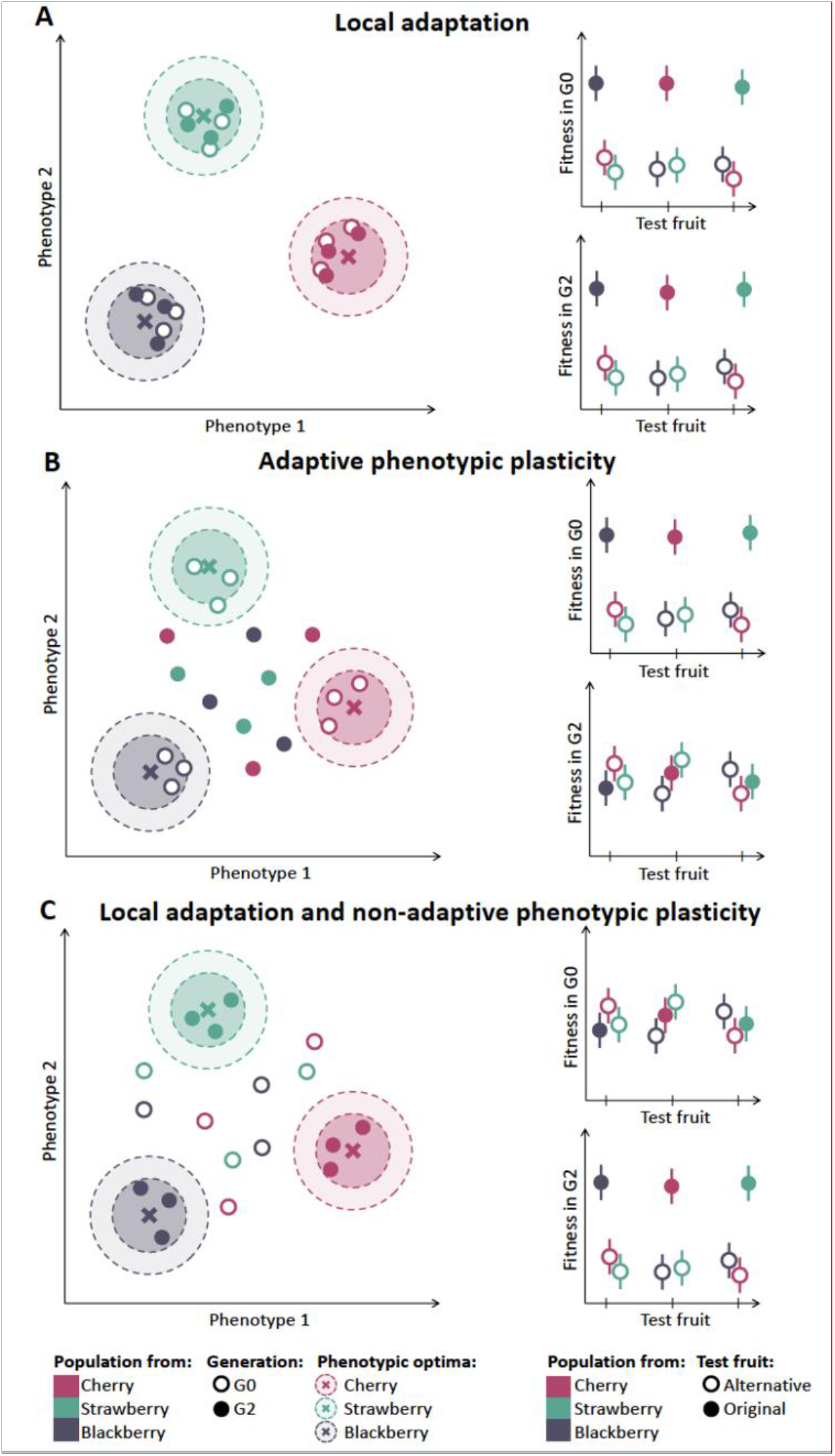
Theoretical predictions for (A) local adaptation, (B) adaptive phenotypic plasticity, and (C) local adaptation with non-adaptive phenotypic plasticity. In the left panels, the two-dimensional fitness landscapes are represented. For each fruit, the position of the phenotypic optimum providing maximal fitness is represented by a cross. The positions of populations in G0 are represented by open and G2 by closed circles. The right panels provide the fitness measures in G0 or G2 in the original and in alternative fruits (closed and open circles, respectively).

Phytophagous insects are valuable biological models to investigate the relative contribution of genetic and plastic responses to varying natural environmental conditions. The host plants of these insects represent heterogeneous resources that vary spatially and temporally in both availability and quality. Oviposition preference for and offspring performance on a host plant are important fitness components that shape host use and specialization in phytophagous insects (Jaenike, 1978; Ravigné *et al*., 2009) and can either be plastic (e.g., learned preference for the habitat encountered as offspring; Dury & Wade, 2020) or genetic (e.g., preference of individuals for the environment they are the best adapted to; Ravigné *et al*., 2009). When hosts co-occur and are present in the same location at the same time, phytophagous insects may maximize their fitness by choosing the best host (e.g., best quality; Jaenike, 1978). However, the host distribution of most phytophagous insects is often spatially or temporally heterogeneous (Denno & Dingle, 1981). Under this scenario, we expect the preference for the host of origin to evolve, as searching for a better host may involves numerous costs (e.g., increased predation risk, increased energetic expenditure, increased risk of inbreeding due to low population size; Papaj & Prokopy, 1989; Davis & Stamps, 2004). In the present study, we focused on the spotted wing drosophila, *Drosophila suzukii*, a generalist crop pest with a broad range of host fruits (Lee *et al*., 2011). This invasive insect species persists throughout the growing season on a succession of different host species as they become available (Kenis *et al*., 2016). Hence, wild populations of *D. suzukii* evolve in a changing environment. Given the seasonality of the fruits and the development time of the insect (Burrack *et al*., 2013; Poyet *et al*., 2015; Aly, 2018), they likely spend only three or four generations on a given host fruit. In contrast to most other generalist *Drosophila* species, the hosts used by *D. suzukii* are well known (Lee *et al*., 2011; Walsh *et al*., 2011). Hence, wild *D. suzukii* meta-populations represent tractable systems to experimentally estimate the relative contributions of genetic and plastic effects to phenotypic adaptation.

To characterize phenotypic adaptation in this species, we sampled natural populations on different fruits and performed a reciprocal common environment experiment over multiple generations in the laboratory. We tested for a pattern of higher performance on or oviposition preference for the fruit they were sampled from, relative to alternative fruits. To do this, we developed a statistical method to estimate the relative contributions of genetic and plastic effects in phenotypic adaptation. Additionally, we examined if oviposition preference and offspring performance traits were correlated, which could speed their co-evolution (Jaenike, 1978).

## Material and methods

### Population sampling and laboratory maintenance

We investigated the adaptation of natural *D. suzukii* populations to three host plants: *Prunus avium* (cherry), *Fragaria × ananassa* Duch (strawberry) and *Rubus fruticosus* (blackberry). We chose these fruit species to represent agronomically important crops (cherry, strawberry) and a wild host fruit considered to be an important reservoir (blackberry; Lee *et al*., 2011; Poyet *et al*., 2015). In the Northern hemisphere, most varieties of cherry and strawberry are available mainly from the end of the spring to the end of the summer, while blackberry is mostly available from the middle of the summer to the middle of the fall. These three species cover the main active period of *D. suzukii* in the Northern hemisphere (i.e., from May to October; Walsh *et al*., 2011).

Between May 2018 and October 2018, we collected fruits from a total of 47 sites in the South of France (Fig. 2, Table S6). The sampled sites included 20, 12, and 15 sites for cherry, strawberry, and blackberry, respectively. For each host plant, fruits were collected towards the end of the production season of the plant to maximize the number of generations over which flies could adapt to their hosts. Recent Capture Mark Recapture studies show that dispersal abilities of *D. suzukii* are generally low relative to the distance between our sampling sites (Tait *et al*., 2018, Tait *et al*., 2019, Vacas *et al*., 2019). Populations hence likely evolved on the sampled host for several generations. In addition, if we happened to sample offspring of recent immigrants, that would merely weaken our ability to detect patterns of local adaptation, and thus any such bias would be in a conservative direction. Field-collected fruits of each population were brought back to the laboratory and kept at 21°C, 65% relative humidity and an 18:6 (L:D) hour cycle in large plastic cages (volume ∼90L) until adult *D. suzukii* flies emerged. This sampling scheme was used consistently throughout the entire experiment. Only the sites from which more than 150 individuals emerged were included in the experiments, leaving a total of 25 sites out of the original 47 (9 cherry, 3 strawberry, and 13 blackberry sites).

**Figure 2.**
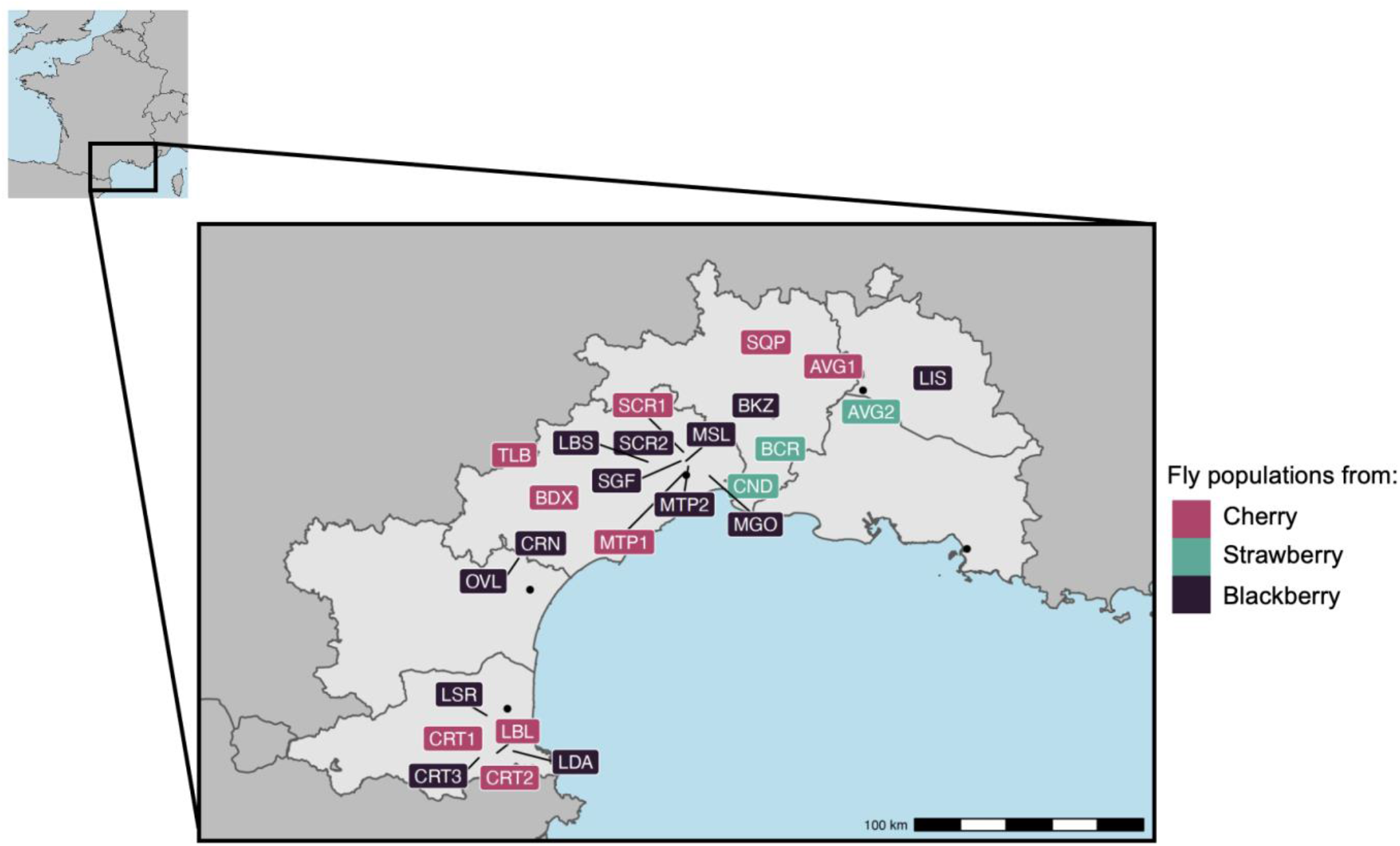
Geographic locations of the 25 sample sites where collected fruits (cherry, strawberry or blackberry) yielded enough *D. suzukii* adults in the lab to be included in this study. See Materials and Methods and Table S6 for details.

### Reciprocal common environment experiment

We measured female preference, fecundity and offspring performance of field-collected flies and lab reared flies on purees made from original fruits (Fig. 3).

**Figure 3.**
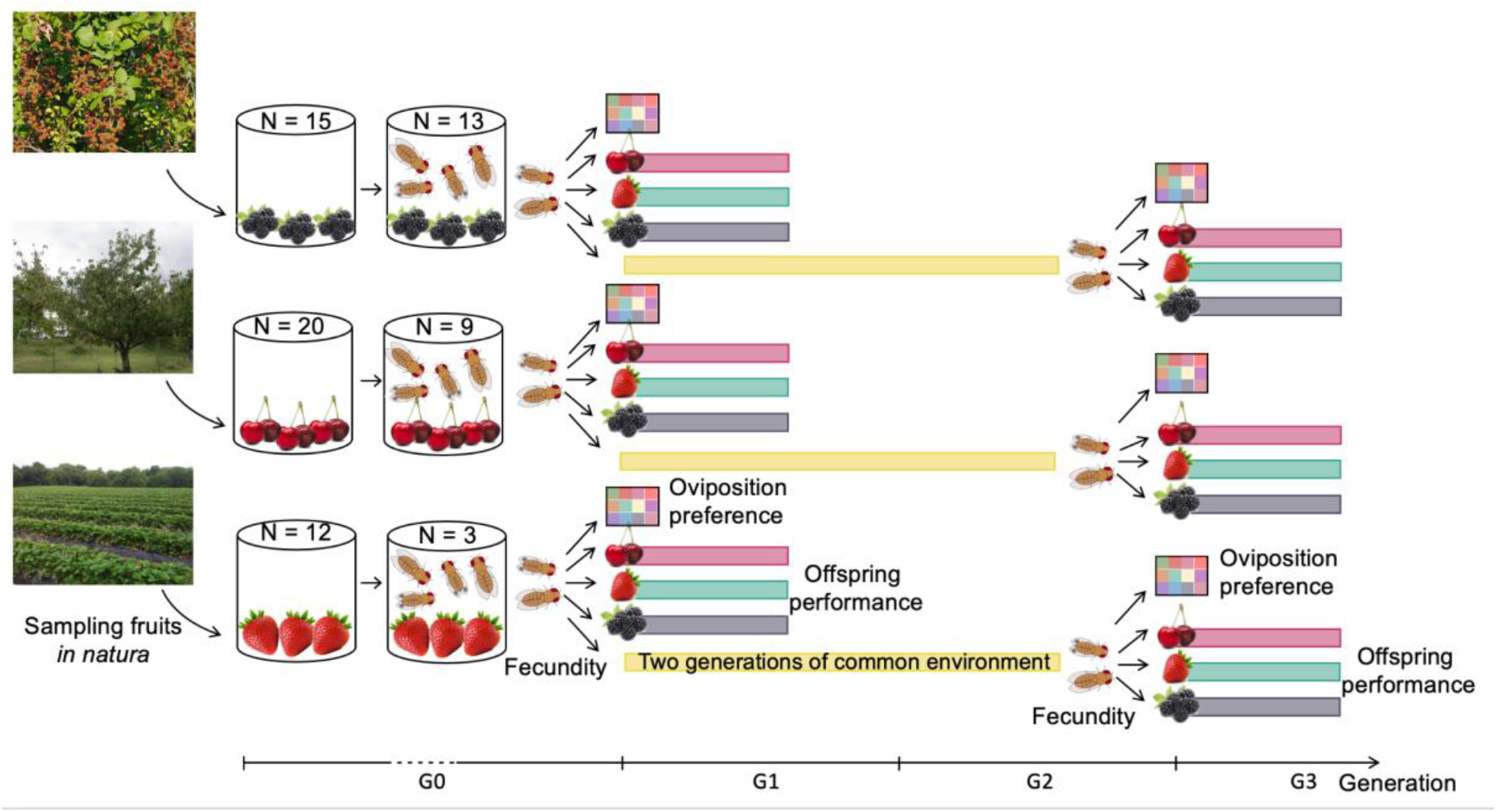
Graphical representation of the experimental design. A standard laboratory fly food medium instead of a fruit puree medium was used for the two generations in a common environment. To test for potential temporal variability in experimental conditions, a control inbred *D. suzukii* line (WT3) was assayed together with each studied population (see main text for details).

To maintain a constant quality of all fruits throughout the experiment (experimental measurements took place over several months), we prepared media using frozen purees of each of the three fruits rather than whole fruits. Batches of media were made with 600 ml of each fruit puree, supplemented with 400 ml of sterile deionized water, 1% (wt/vol) of agar, 6% of inactive malted brewer’s yeast, and 1.5% of yeast extract. We then added an antimicrobial solution, which consisted of 6 ml of 99% propionic acid, 10 ml of 96% ethanol and 0.1% of methylparaben sodium salt. For the common environment, we used a standard laboratory fly food (i.e., “German food”; Backhaus *et al*., 1984) including 1000 ml of sterile deionized water, 1% (wt/vol) of agar, 6% of dextrose, 3% sucrose, 8% of inactive malted brewer’s yeast, 2% of yeast extract, 2% of peptone, 0.05% of MgSO_4_, and 0.05% of CaCl_2_, and 16 ml antimicrobial solution (6 ml of 99% propionic acid, 10 ml 96% of ethanol and 0.1% of methylparaben sodium salt). See Olazcuaga *et al*., 2019 for product references.

We collected adult flies from the field-collected fruits after the peak of emergence (usually within four days after the first adults emerged). We kept only *D. suzukii* individuals and discarded other *Drosophila* species (Hauser, 2011). We placed adults (hereafter referred to as “G0”) in cages with organic commercial fruits corresponding to the fruits from which the population originated and allowed them to mature for seven days (the absence of prior *D. suzukii* infestation of these fruits was assessed by rigorous visual inspection). We performed reciprocal common environment experiments in the laboratory using (*i*) seven-day-old (+/- two days) G0 adults that emerged from field-collected fruits and (*ii*) seven-day-old (+/- one day) G2 adults obtained after two generations of maintenance in a common environment (i.e., “German food”; Backhaus *et al*., 1984). Unlike an experiment where all populations would grow under controlled conditions in all test environments and then be measured in each test environment (i.e., a “two-way” experimental design), the two-generation experimental design we used enabled us to detect potential adaptive phenotypic plasticity in responses to environmental cues that are independent of the fruit itself (e.g., the photoperiod, the temperature, etc.). Using a two-way experimental design by rearing laboratory flies on different fruits for two generations under laboratory conditions could have decreased our power to detect adaptive plasticity. For each population, we estimated female preference and offspring performance in artificial fruit media (hereafter test fruit) composed of either the fruit from which the population was sampled (hereafter original fruit) or two other alternative fruits. We measured oviposition preference and fecundity on each test fruit (see below) in G0 and G2, as proxies for female preference and we measured egg-to-adult survival in G1 and G3, as a proxy for offspring performance.

Oviposition preference was measured as the number of eggs laid on different fruits by 20 females in 24 hours in an arena that contained the media of the three above-mentioned host fruits plus 9 other test fruit media (apricot, blackcurrant, cranberry, fig, grape, kiwi, raspberry, rose hips, and tomato) distributed randomly into 12 compartments (see Fig. S2 in Olazcuaga *et al*., 2019). Hence, these arenas contained a wide selection of *D. suzukii* hosts that are present in the region from which wild populations were collected, expanding the inference space in which we measure oviposition preference. The detection of the oviposition preference for a fruit in these arenas is conservative, since the preference in a choice environment of 12 fruits would likely be present in a choice environment containing only three of these fruits. In addition, this experimental design has been used in other studies (e.g., Olazcuaga *et al*., 2019), and using this same design facilitates comparison of the oviposition preference of these wild populations with other wild or experimental populations. We also used a no-choice assay to measure the oviposition response of females to individual fruits (hereafter fecundity), as the number of eggs laid when placing 20 adults for 24 hours in a vial with one of the three fruits. Eggs were counted under dissecting microscopes. For G0, we used on average 7.8 arenas per population to estimate oviposition preference and an average of 7.5 vials per population for the other traits. For G2, we used 7.9 arenas and 8.0 vials, respectively.

Offspring performance was measured on each fruit as egg-to-adult survival. We quantified egg-to-adult survival as the proportion of eggs in the no-choice assay that resulted in the emergence of adults 16 days after oviposition (more than 95% of individuals emerge within 16 days on cherry, strawberry, or blackberry media; Fig. 4 in Olazcuaga *et al*., 2019). It is worth noting that because *D. suzukii* oviposits directly into the media, removing eggs to control density was too time consuming given the size of the experiment. Furthermore, manipulation of the eggs reduces survival (Schou, 2013). We therefore accounted for egg density afterwards during statistical analysis (see statistical methods section) rather than during the experiment. Egg density did not vary consistently across test or original fruits, and thus accounting for them during statistical analysis should not lead to bias. More precisely, 77.7% of the variance in the egg density was explained by the variation across tubes, while only 0.6% of the variance was explained by variation among test fruits and 21.7% by variation among populations.

**Figure 4.**
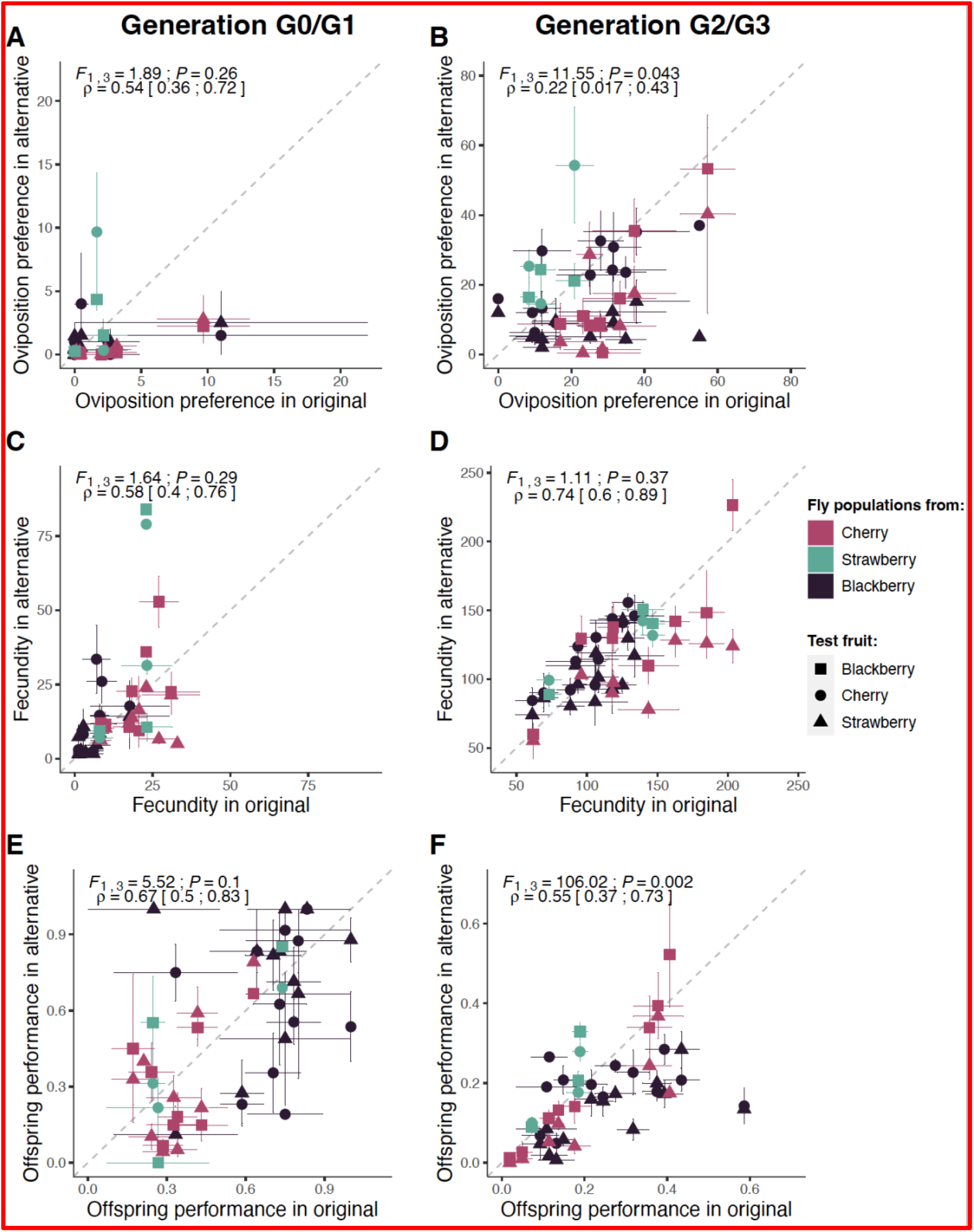
Relationship between original and alternative fruits for (A-B) oviposition preference (number of eggs laid by 20 females in 24 hours in the original or alternative fruit media, in a choice environment), (C-D) fecundity (number of eggs laid by 20 females in 24 hours in the original or alternative fruit media, in a no-choice environment), and (E-F) offspring performance (egg-to-adult survival) in generations G0 or G1 (left panels) and G2 or G3 (right panels). The term original in the x-axis legend indicates that the trait is measured in the same fruit from which populations were collected, while the term alternative in the y-axis legend indicates measurements on other fruits. The mean of each population in each test fruit is represented by a symbol whose shape depends on the test fruit (fruit medium) and color depends upon the fruit from which the population was collected *in natura*. For each population, the two means measured in the two alternative fruits are associated with the same and unique mean in the original fruit. Local adaptation corresponds to the cases where the means measured in the two alternative fruits are both located below the line of symmetry, due to a higher mean trait value in the original fruit. Error bars represent the standard error of mean estimates. Results (*F*-statistics and *P*-values) from the SA method detailed in Blanquart *et al*., (2013), as well as the weighted Pearson’s correlation coefficient (ρ) with their 95% confidence interval, are given above each panel.

Our field sampling design followed the seasonality of each fruit. As a result, reciprocal common environment experiments of each population were performed at different dates. To test for potential temporal variability in experimental conditions, we used an inbred *D. suzukii* line (*WT3*; Chiu *et al*., 2013) as a control. Together with each wild population, we measured oviposition preference, fecundity, and offspring performance in this control population. Analyses of each of the three traits of interest showed that temporal variation was of the same order of magnitude as the variation within a given date (Appendix S1).

### Statistical methods

The analyses of oviposition preference and fecundity (log-transformed number of eggs; Miller, 1997) and offspring performance (arc-sine-transformed egg-to-adult survival; McDonald, 2014) were performed using R statistical software (R Core Team, 2014). For offspring performance, 51 vials had more adults than the number of eggs counted (50 of 541 vials for G0 and 1 of 602 for G2, with an average egg-to-adult survival observed of 2.44) due to a difficulty to count the eggs in the vials with very few eggs (average number of eggs of these 51 vials of 3.65). We assumed an egg-to-adult survival of 1 in the analyses of the offspring performance for these vials. Doing so is not expected to introduce a bias in the data analysis because such vials were equally represented in all treatments (17, 16 and 18 observations in cherry, strawberry and cranberry vials, respectively). In agreement with this, processing our egg-to-adult survival statistical analyses provided the same qualitative results when these 51 observations were removed (results not shown).

#### Testing for trait differences between original and alternative fruits

For each trait and each generation separately, we tested whether populations had a higher preference for or performance in their original fruit than in alternative fruits. First, to visualize the patterns, we plotted the average trait in the medium corresponding to alternative fruits against the same average trait in the medium corresponding to the original fruit for each population. In addition, we computed the weighted correlation coefficient between the average trait in alternative fruits and their average in the original fruits, using the total number of vials for each population as weight and estimated its 95% confidence interval using the *sjstats* package (Lüdecke, 2018).

Second, we tested for differences in oviposition preference, fecundity and offspring performance of populations between original and alternative fruits using the SA method (where S stands for sympatric and A stands for allopatric) detailed in Blanquart *et al*., (2013). Unlike the ‘home-away’ and ‘local-foreign’ methods (Kawecki & Ebert, 2004), the statistical power of the SA method does not decline with increasing sample size (number of populations; Blanquart *et al*., 2013). Most notably, the SA method controls for potential variation in overall quality of the test environments, a feature previously reported for hosts of *D. suzukii* (Bing *et al*., 2018; Olazcuaga *et al*., 2019). Additionally, the SA method controls for confounding factors due to quality variations among populations such as differences in inbreeding depression (Keller, 2002) or differences in the prevalence of parasites or endosymbionts transmitted across generations (Fry *et al*., 2004; Merçot & Charlat, 2004). Finally, using an ANOVA model rather than a Linear Mixed Model, decreases the rate of false positive detection of local adaptation (Appendix E in Blanquart *et al*., 2013).

The SA method we used is detailed in Appendix S2. Briefly, it consists of an ANOVA-based approach to fit a model that includes population (accounting for both intrinsic genetic differences and environmental effects induced by differences among original fruits in G0/G1), test fruits (accounting for differences in dietary quality among test fruits) and an interaction between original fruits and test fruits. The model also includes a factor, called *SA*, which indicates whether the test fruit is the original (sympatric) fruit or an alternative (allopatric) fruit. The significance of the *SA* effect is tested using an *F-*test where the ratio evaluates the variation due to *SA* over the total variation minus that due to habitat and population quality (Blanquart *et al*., 2013, Appendix D). Positive values of *SA* indicate a pattern of local adaptation, while negative values indicate local maladaptation.

#### A new method to detect and quantify local adaptation and adaptive phenotypic plasticity simultaneously

To evaluate the relative contributions to preference and performance of local adaptation and adaptive phenotypic plasticity, it was necessary to analyze the data across the G0/G1 and G2/G3 generations. When measured on adults that emerged from field-collected fruits (G0/G1), phenotypic differences among populations can be driven both by genetic differences among populations (potentially including local adaptation) and by plastic responses to the natural environment (potentially including adaptive phenotypic plasticity). When measured on adults after two generations of common environment (G2/G3), phenotypic differences among populations are likely to have a genetic basis, as potential maternal and grand-maternal environmental effects are standardized across populations. Although transgenerational plasticity and other non-genetic factors such as vertically transmitted symbionts can still be present, we use the term “genetic effects” hereafter for the sake of brevity. To evaluate the relative contributions to preference and performance of local adaptation and adaptive phenotypic plasticity, we developed new approaches to visualize and test for genetic and plastic effects. Briefly, by using a custom model that compares the results of the two trials, the G0/G1 generation and the G2/G3 generation, we estimated the variance components and effects that could be attributed to local adaptation and adaptive phenotypic plasticity.

To visually illustrate genetic and plastic effects in preference, we estimated the mean of genetic effects based on preference data from G2 and the mean of plastic effects as the difference in preference between G0 and G2, after controlling for other sources of variation, including variation in quality among test fruits and among populations, as well as variation among arenas for oviposition preference or among vials with different egg densities for offspring performance (Appendix S3). Similarly, for performance, we estimated the mean genetic effects from G3, and the mean of plastic effects as the difference between G1 and G3.

To statistically test for genetic and plastic effects, we modified the SA method detailed in Blanquart *et al*., (2013) to test whether oviposition preference, fecundity and offspring performance of populations was on average different in the original fruits and in alternative fruits and whether this effect was either genetic and present in both generations or was plastic and present in the first generation (G0 or G1), but not the second generation (G2 or G3). To this end, we derived two different test statistics for genetic and plastic effects respectively (*F*_*genetic*_ and *F*_*plastic*_; see Appendix S4 for details). We performed computer simulations to assess the general properties of our method. We found that both genetic and plastic effects can be reliably detected using our method under both balanced and unbalanced experimental designs, with a false positive rate for the detection of genetic and plastic effects <3% (Fig. S5; Tables S3, S4 and S5). We also found that for unbalanced designs (as in the present study) using an ANOVA model, rather than a Linear Mixed Models substantially increased the power of detecting local adaptation and adaptive phenotypic plasticity (Appendix S4).

For oviposition preference, we fitted the following ANOVA model on the log-number of eggs, *y*_*ijklm*_:

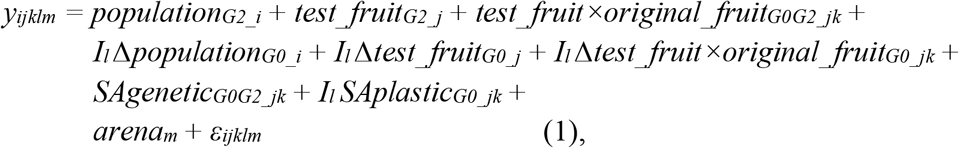

where *I*_*l*_ is an indicator variable with *I*_*l*_ =1 when generation *l* is G0 and *I*_*l*_ =0 otherwise, and fixed effects included the effect of the *i*th population estimated in G2 (*population*_*G2_i*_ with *i*=1,…,25) to account for differences in quality among populations, the *j*th fruit media estimated in G2 (*test_fruit*_*G2_j*_ with *j*=1,…,3 for blackberry, cherry and strawberry, respectively) to account for differences in quality among test fruits, the interaction between the *j*th test fruit and the *k*th original fruit that is observed in both G0 and G2 (*test_fruit×original_fruit*_*G0G2_jk*_), and the plastic difference of these effects in G0 relative to the same effects in G2 (*Δpopulation*_*G0_i*_, *Δtest_fruit*_*G0_j*_, *Δtest_fruit×original_fruit*_*G0_jk*_ respectively). Fixed effects also included two SA effects, which indicated whether the test fruit was the original, sympatric, fruit or was an alternative, allopatric fruit in either G0 or G2 (*SAgenetic*_*G0G2_jk*_) and in G0 (*SAplastic*_*G0_jk*_). Hence, *SAgenetic*_*G0G2_jk*_ measures local adaptation in both G0 and G2 and *SAplastic*_*G0_jk*_ measures adaptive phenotypic plasticity in G0 (i.e., field-collected flies). The model also included the fixed effect of the *m*th arena in G0 and G2 (*arena*_*m*_ with *m*=1,…,179 in G0 and *m*=180,…, 369 in G2) to account for differences among arenas and a random error to account for the variation among observations from the same arena (*ε*_*ijklm*_, normally distributed with a mean of zero and variance *σ*^*2*^_*res*_). As with the original SA method, this new method detects if there is a significant difference when the test fruit is the original fruit *vs*. an alternative fruit. A significant difference in the associated *F*-test with a positive estimate of *SAgenetic*_*G0G2*_ and *SAplastic*_*G0*_ provides support in favor of local adaptation and adaptive phenotypic plasticity, respectively. We either analyzed the entire dataset (12 fruits) or focused on a smaller dataset with the three fruits of interest (cherry, strawberry, blackberry). Results were similar in both analyses, so we only present the results with only the three fruits of interest.

For fecundity, we fitted the same model but without the arena effect. For offspring performance, we fitted the same model with G1 and G3. Offspring performance increases with egg density and levels off at high egg densities (Fig. S12). To account for egg density in the statistical analysis of offspring performance, we replaced the arena effect by the log-transformed initial number of eggs (*eggs*_*m*_).

To estimate the level of local adaptation, we designed a new variable *φ*_*genetic*_ based on the proportion of variance of the interaction *test_fruit×original_fruit*_*G0G2_jk*_ explained by *SAgenetic*_*G0G2_jk*_:

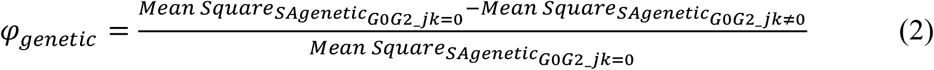

Similarly, the level of adaptive phenotypic plasticity *φ*_*plastic*_ was estimated based on the proportion of variance of the interaction *test_fruit×original_fruit*_*G0_jk*_ explained by *SAplastic*_*G0_jk*_:

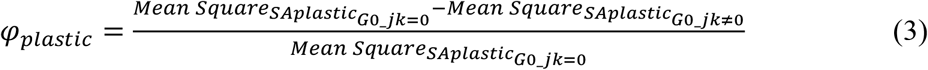

Because we did not detect any significant adaptive phenotypic plasticity (see results), we retrospectively performed a simulation-based analysis to estimate our power to detect the presence of adaptive phenotypic plasticity with our experimental design (Appendix S5).

#### Testing for correlations between generation and between fitness related traits

We tested for two sets of correlations. First, we evaluated whether there is a correlation in oviposition preference, fecundity or offspring performance of populations between generations (Appendix S6). Second, we examined whether the evolution of preference and performance traits are independent or rather if they coevolve. Specifically, we tested for a significant correlation between oviposition preference and offspring performance between G0 and G1, and between G2 and G3 (Appendix S7). In each of these two cases, we estimated correlations using the total number of vials for each population as a weight and estimating its 95% confidence interval using the *sjstats* package (Lüdecke, 2018).

## Results

### Test for trait differences between original and alternative fruits

Across the three traits of interest (oviposition preference, fecundity and offspring performance) and for each generation separately, populations with higher trait values in their original fruit generally had higher trait values in alternative fruits (Fig. 4). Indeed, the correlation coefficients between traits in original and alternative fruits were positive and significantly different from zero for all the traits and generations (Fig. 4). These positive correlations indicate variation in intrinsic quality among populations within and among fruits and, most importantly, emphasizes the importance of statistically accounting for this variation when testing for local adaptation and adaptive phenotypic plasticity (Appendix S2). For oviposition preference, these correlations among populations were higher in G0 than in G2 for oviposition preference (ρ_G0_ = 0.54 and ρ_G2_ = 0.22; Fig. 4A vs. 4B). In contrast, for fecundity, the correlations among populations were weaker, with more variation observed in G0 than in G2 (ρ_G0_ = 0.58 and ρ_G2_ = 0.74; Fig. 4C vs. 4D), suggesting greater phenotypic plasticity in this trait in flies collected from the field. Finally, these correlations among populations were higher in G1 than in G3 for offspring performance (ρ_G1_ = 0.67 and ρ_G3_ = 0.55; Fig. 4E vs. 4D).

In G0 and G1, the values of the three traits of interest did not significantly differ between original and alternative fruits (Fig. 4A, C, E, Fig. S5A, C, E). In G2 and G3, most of the populations preferred to oviposit and had higher offspring performance in their original fruit than in alternative fruit media (i.e., most populations are under the dashed line of symmetry in Fig. 4B, F). This pattern was particularly marked for offspring performance (*F*_1,3_ = 106.02, *P-value* = 0.002; Fig. S5F), but more tenuous for oviposition preference (*F*_1,3_ = 11.55, *P-value* = 0.043; Fig. S5B). In contrast, no evidence for higher fecundity on the original fruit than on alternative fruit media was found (*F*_1,3_ = 1.11, *P-value* = 0.37; Fig. 4D; Fig. S5D).

### Relative importance of local adaptation and adaptive phenotypic plasticity

When analyzing G0 and G2 data together (Fig. 5), our new statistical method showed that the higher preference for their original fruit media had a significant genetic origin (*P-value* = 0.037 with a positive estimate of *SAgenetic*_*G0G2*_; Table 1). Overall, local adaptation (*SAgenetic*_*G0G2*_) explained 74.64% of the variance of the genetic interaction between the test and original fruits in G0 and G2. While phenotypic plasticity was in the maladaptive direction (negative estimate of *SAplastic*_*G0*_; Table 1), it was not significantly different from zero (*P-value* = 0.16).

**Table 1:** Estimates for *SAgenetic* and *SAplastic* for oviposition preference, fecundity, and offspring performance. The *SAgenetic* and *SAplastic* estimates are presented with their standard error (SE_*SAgenetic*_ and *SE*_*SAplastic*_ respectively). The *Fgenetic* and *Fplastic* on *SAgenetic* and *SAplastic* and corresponding *P-values* are also indicated.

|  | Oviposition<br>preference | Fecundity | Offspring<br>performance |
| --- | --- | --- | --- |
| $\Phi_{genetic}$ | 74.64% | 18.67% | 80.90% |
| $\Phi_{plastic}$ | 38.81% | -0.27% | -5.01% |
| $F_{genetic}$ | 12.77 | 1.92 | 17.94 |
| $P$-value | 0.037 | 0.26 | 0.024 |
| $SA_{genetic}$ | 0.384 | -0.058 | 0.042 |
| $SE_{SA_{genetic}}$ | 0.08 | 0.063 | 0.028 |
| $F_{plastic}$ | 3.54 | 0.99 | 0.81 |
| $P$-value | 0.16 | 0.39 | 0.43 |
| $SA_{plastic}$ | -0.266 | -0.132 | 0.043 |
| $SE_{SA_{plastic}}$ | 0.116 | 0.099 | 0.044 |

**Figure 5.**
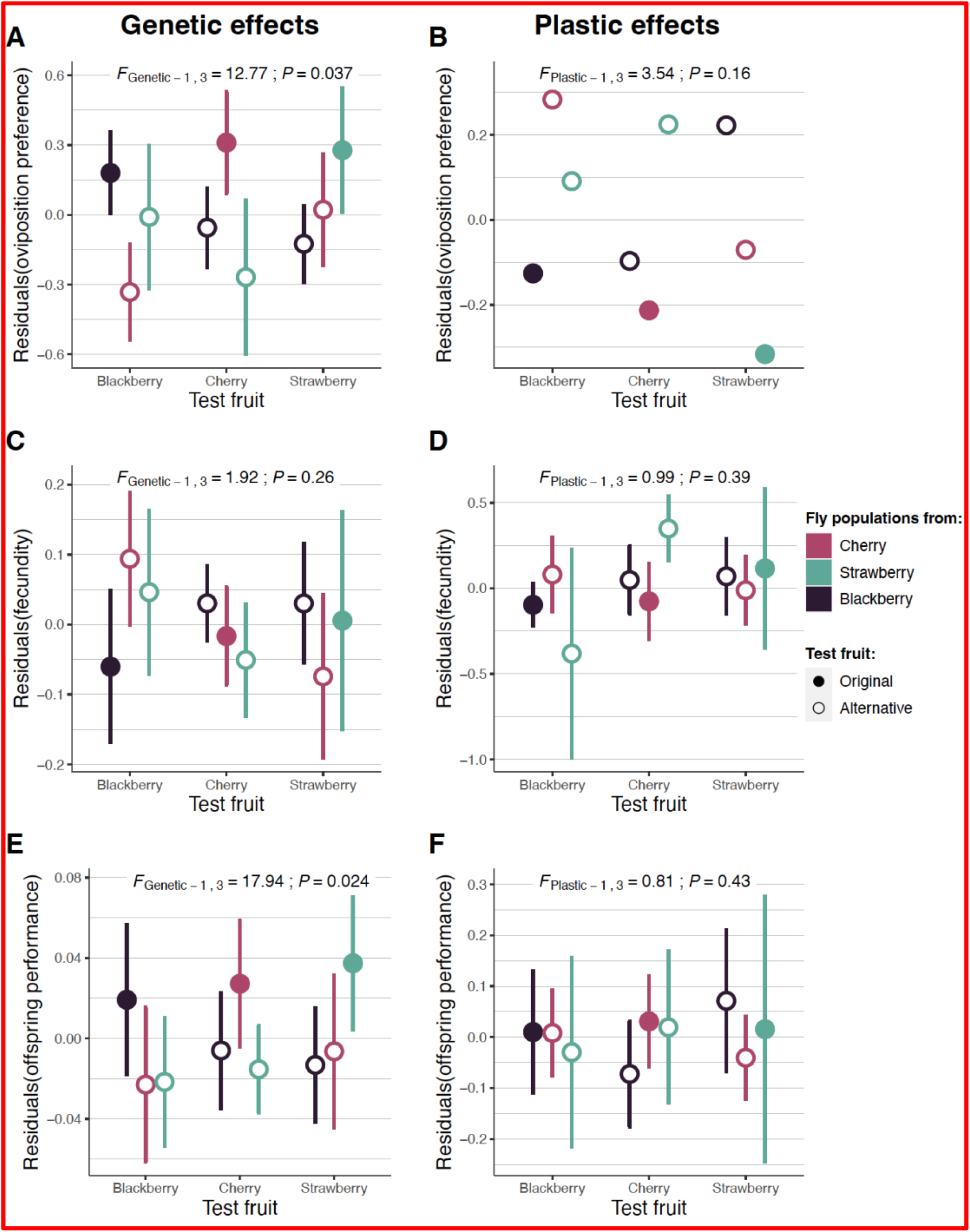
Genetic (left panels) and plastic (right panels) effects across *D. suzukii* natural populations for (A-B) oviposition preference, (C-D) fecundity, and (E-F) offspring performance for each combination of original fruit and test fruit. Genetic effects (i.e., testing for local adaptation) correspond to each trait value in G2 or G3, while plastic effects (i.e., testing for adaptive phenotypic plasticity) correspond to the differences between each trait value in G0 or G1 and their respective trait value in G2 or G3. For both genetic and plastic effects, we ran models to account for other sources of variation, including variation in quality among test fruits and among populations, as well as variation among arenas for oviposition preference or among vials with different egg densities for offspring performance. The residuals from these models are represented on the y-axis. Results (*F*-statistics and *P*-values) for the test of adaptive phenotypic plasticity or local adaptation are given above each panel. Error bars represent 95% confidence intervals. For plastic effects of oviposition preference, 95% confidence intervals could not be computed due to a higher variance in G2 than in G0.

For fecundity, the absence of pattern in G0 and G2 when analyzed independently (Fig. S5) was confirmed, as no significant genetic or plastic effects were detected when analyzing G0 and G2 data together (Fig. 5C-D, *P-value* = 0.26 and *P-value* = 0.39 respectively; Table 1).

The higher performance of the flies on their original fruit media (Fig. 5F) had a significant genetic origin (*P-value* = 0.024, with a positive estimate of *SAgenetic*_*G1G3*_; Table 1). Overall, local adaptation (*SAgenetic*_*G1G3*_) explained 80.9% of the variance of the genetic interaction between the test and original fruits in both G1 and G3. While phenotypic plasticity was in the adaptive direction for this trait (positive estimate of *SAplastic*_*G1*_; Table 1), it was not significantly different from zero (*P-value* = 0.43).

### Testing for correlations between female preference and offspring performance

We tested for a correlation between oviposition preference and offspring performance, after controlling for other sources of variation, including variation in quality among test fruits and among populations, as well as variation among arenas for oviposition preference or among vials with different egg densities for offspring performance (see Appendix S7 for details). We found that oviposition preference in G0 was positively correlated with offspring performance in G1, and oviposition preference in G2 was positively correlated with offspring performance in G3 (Fig. S9). The correlation coefficient was significantly different from zero for the G2/G3 generation test, whereas the 95% confidence intervals of the correlation coefficient included zero for the G0/G1 generation test.

## Discussion

We used natural populations of *D. suzukii* to evaluate the roles of local adaptation and adaptive phenotypic plasticity in driving fitness adjustment to changes in the environment occurring over an intermediate pace of 3-4 generations. We focused on three key traits: oviposition preference, fecundity and offspring performance. We found local adaptation for oviposition preference and offspring performance, but not for fecundity. Females preferred their original fruits, and their offspring performed better on those fruits. We did not find any signature of adaptive phenotypic plasticity for any of the three traits. If adaptive phenotypic plasticity plays a role in the fit to the environment, that role is small enough to be masked by the effects of non-adaptive plasticity and local adaptation.

### Strong variation in quality among populations in all tested fruits

For each of the three traits, *D. suzukii* populations displayed consistent correlations between the original and test fruits. Some populations had the stronger preference or performance across tested fruits, highlighting the high variation in intrinsic quality among populations. *D. suzukii* population quality may be related to various ecological and historical factors, as well as their genetic background, with some populations having, for instance, low genetic load and others having higher genetic load. The level of inbreeding depression, which depends on the level of genetic load, is indeed known to vary among wild populations and influences life history traits (Keller and Waller, 2002).

### The higher preference and performance in original fruit media is driven by local adaptation rather than adaptive phenotypic plasticity

We found that the stronger oviposition preference for and higher performance in the original fruit than in alternative fruits was due to genetic and not plastic responses. Our research is relevant to current understanding of how the tempo of environmental change over time and space (i.e., environmental grain) shapes the way organisms improve the match between their phenotypic traits and their local environment. Based on the period of availability of each host fruit, on average temperatures in the studied areas and the generation time estimates in the laboratory (Poyet *et al*., 2015; Olazcuaga *et al*., 2019), our results indicate that local adaptation (and not just adaptation) can evolve in less than four generations in large diverse natural populations of *D. suzukii*. This indicates that the evolutionary process of local adaptation, including lower relative fitness in alternative fruits, can be more rapid than traditionally thought (e.g., 10 generations in natural populations of *D. melanogaster*; Bergland *et al*., 2014). In agreement with this, rapid adaptation over only a few generations have been recently documented in populations of *D. melanogaster* evolving in semi-natural conditions (Rudman *et al*., 2021) and natural conditions (Machado *et al*., 2021).

It is likely that at the end of the season of a given fruit, the *D. suzukii* populations established on this fruit will switch to another fruit resource located in the neighborhood since diapause has never been observed for this species between spring and the end of summer (Walsh *et al*., 2011). Supporting this interpretation is the fact that genetic differentiation among populations sampled on different fruits is weak (unpublished data). Therefore, because populations have to switch to a new fruit regularly due to the seasonality of fruits, the rapid and dynamic process of local adaptation in *D. suzukii* seems to be transient. Transient and rapid local adaptation has also been found in experimental populations of another crop pest species *Tetranychus urticae* maintained in the lab in a temporally heterogeneous environment with two host plants (cucumber or pepper) in succession (Bisschop *et al*., 2019). At the genome level, the seasonality of host fruits might result in balancing selection at adaptive loci and thus help to maintain high standing genetic variation in natural *D. suzukii* populations at and around those loci (Gloss *et al*., 2013).

The rapid genetic adaptation we observed could be favored by *D. suzukii* populations having a large effective size and by strong and divergent selective pressure exerted by seasonal fruits. The population size of *D. suzukii* in each of the studied sites is likely to be large (on the order of 10^4^ adults according to estimates from American cherry orchards; Tochen *et al*., 2014; Wiman *et al*., 2014) as suggested by the high levels of genetic diversity observed in *D. suzukii* populations sampled in the South of France (Fraimout *et al*., 2017; Olazcuaga *et al*., 2020). Differences in chemical composition among fruits likely result in strong pressures on traits such as offspring performance, as evidenced by a recent experimental evolution study (Olazcuaga *et al*., 2021).

We did not detect a correlation between trait values measured in generations G0 and G2 for preference traits or in G1 and G3 for performance for a given population. This finding can be explained by the presence of a substantial amount of non-adaptive plasticity in G0 (Fig. 1C), which masked the genetic effects responsible for phenotypic patterns of local adaptation. In addition, we did not detect adaptive phenotypic plasticity although we had enough statistical power to detect an effect of the same magnitude as local adaptation. The failure to detect adaptive phenotypic plasticity may seem surprising at first sight because *Drosophila* species, including *D. suzukii*, usually excel at using cues to optimize their oviposition behavior (Jaenike, 1983; Papaj & Rausher, 1983; Little *et al*., 2020). The absence of evidence for adaptive phenotypic plasticity suggests that in the case of *D. suzukii* the changes in plant hosts are not frequent enough for plasticity to be strongly beneficial. In addition, the flies were sampled in an agricultural region in France that tends to exhibit monocultures of crops rather than polycultures, which might also contribute to a lack of plasticity, as different crop fruits are rarely available at the same time within the same area.

### Relationships between female oviposition preference and offspring performance

We found that oviposition preference in generation G2 was positively and significantly correlated with offspring performance in generation G3. Previous studies that tested for this correlation in *D. suzukii* have shown mixed results (Poyet *et al*., 2015; Olazcuaga *et al*., 2019; Shu *et al*., 2021b). Our results, based on natural populations of *D. suzukii*, suggest that selection pressures on oviposition preference might drive adaptation to host fruits and could speed up local adaptation. This is consistent with previous theoretical and experimental studies showing that selective pressures on oviposition preference could feedback positively on the evolution of performance for local hosts (Whitlock, 1996; Berlocher & Feder, 2002; Via & Hawthorne, 2002). In particular, if preference and performance traits are genetically linked, strong selection pressures on oviposition preference, at least for the three fruits studied here, could lead to the evolution of performance traits (Wood *et al*., 1999; Berner & Thibert-Plante, 2015).

### Limits of our experimental approach

Our experiments do not overcome the common drawback of trying to infer the fitness of natural populations using experiments in the laboratory rather than in natural conditions (Reznick & Ghalambor, 2005). First, the small scale of our set up for studying choice among multiple fruits deviates considerably from the conditions and geographic scales found in natural landscapes and may therefore provide an incomplete picture of the adaptive processes regarding oviposition preference in *D. suzukii*. Second, our experiments did not consider yeasts and bacteria that occur in fruits in the field, and which have been shown to modify the foraging behavior (Shu *et al*., 2021a) as well as the oviposition preference (Bellutti *et al*., 2018) of *D. suzukii* females. Yeasts and bacteria can also affect performance, as microbes represent an important source of protein for the egg-to-adult survival of *D. suzukii* (Lewis & Hamby, 2019; Bing *et al*., 2021). Therefore, if the microbial community differs among cherry, strawberry and blackberry, our results could be at least partly explained by the nutritional effect of a fly microbiome transmitted over generations rather than a *D. suzukii-* based genetic effect. Experiments in *D. melanogaster* have shown that its bacterial community can be transmitted over successive generations in the laboratory (Téfit *et al*., 2018) and that both *Acetobacter* and *Lactobacilli* can grow in media with concentration of antimicrobials similar to the one in our media (Obadia *et al*., 2018). Using axenic individuals could help testing for the role of the microbiome in the patterns we detected, although the absence of microbiota itself could have an important impact on both preference and performance.

## Conclusion

We studied oviposition preference, fecundity and offspring performance in *D. suzukii* to evaluate the relative influence of genetic and plastic effects on the match between phenotypic traits and the local environment, here the host fruit. We found a pattern of local adaptation for oviposition preference and offspring performance, but not for fecundity. We found no evidence of adaptive phenotypic plasticity for all studied traits. Our study hence demonstrates that spatially and temporally variable selection does not prevent the rapid evolution of local adaptation in natural populations over a short number of generations. This dynamic process appears to be transient as populations rapidly adapt to new host fruits as they become available throughout the year. In addition to testing for the potential role of the microbiome in the pattern we detected, a stimulating follow up to our study would be to identify the physiological pathways under selection using genome-environment association study based on populations sampled from different host plants. Such a combination of statistical, molecular, and quantitative approaches would provide useful insights into the genomic and phenotypic responses to divergent selection among host fruits in phytophagous generalist insects. Finally, our new statistical method will likely contribute to the quantification of the relative contribution of local adaptation and adaptive phenotypic plasticity in any other organisms experiencing spatially or temporally variable selection.

## Acknowledgements

We are grateful to V. Ravigné and B. Facon for insightful discussions, P. Audiot, M.P. Chapuis, L. Benoit, N. Vieira, J. Froissard, A. L. Clamens, S. Nidelet and L. Sauné for technical assistance, S. Bonamour for her comments on the manuscript, V. Cazalis and R. Patin for their help with drawing the map, and to the Sicoly cooperative for providing us with some fruit purees. L.O., M.G., J.F. and A.E. were supported by the Languedoc-Roussillon region (France) through the European Union program FEDER FSE IEJ 2014-2020 (project CPADROL) and the INRAE scientific department SPE (AAP-SPE 2016). R.A.H. acknowledges support from the National Science Foundation (DEB-0949619), USDA Agriculture and Food Research Initiative award (2014-67013-21594), Hatch project 1012868, the French Agropolis Fondation (LabEx Agro–Montpellier) through the AAP “International Mobility” (CfP 2015-02), the French programme investissement d’avenir, and the LabEx CEMEB through the AAP “invited scientist 2016”. N.O.R. acknowledges support from the CeMEB LabEx/University of Montpellier (ANR-10-LABX-04-01) and the INRAE scientific department SPE (AAP-SPE 2021PestAdapt).

## Author contributions

Conceptualization and funding acquisition: L.O., J.F., M.G., R.A.H., N.O.R. and A.E.; Statistical analyses and writing: L.O. with inputs of J.F., M.G., R.A.H., N.O.R. and A.E.; Data acquisition and field sampling: L.O., C.D., A.L., J-L.I., R.V., R.G., C.S. and A.E.

## Data accessibility

The data and R scripts for our analyses are available at: https://github.com/nrode/NatPop2021 They will be archived on Dryad upon acceptance of the manuscript.

