## Supplementary materials for "Rapid and transient evolution of local adaptation to seasonal host fruits in an invasive pest fly"

**Short title: Rapid local adaptation in *Drosophila suzukii***

Laure Olazcuaga[1,2], Julien Foucaud[1], Candice Deschamps[1], Anne Loiseau[1], Jean-Loup Claret[1], Romain Vedovato[1], Robin Guilhot[1], Cyril Sevely[3], Mathieu Gautier[1], Ruth A. Hufbauer[1,2], Nicolas O. Rode[1,4] and Arnaud Estoup[1,4]

[1] CBGP, Univ Montpellier, CIRAD, INRAE, Institut Agro, IRD, Montpellier, France
[2] Department of Agricultural Biology and Graduate Degree Program in Ecology, Colorado State University, Fort Collins, Colorado 80523, USA
[3] Chambre d'agriculture de l'Hérault, Lattes, France
[4] These authors contributed equally to this work

**Corresponding authors: Laure Olazcuaga, Nicolas O. Rode and Arnaud Estoup**

**Corresponding author: Laure Olazcuaga**

CBGP - 755 avenue du Campus Agropolis

CS 30016 - 34988 Montferrier sur Lez cedex

France

### Table of content

**Appendix S1**
Table S1
Figure S1-S2

**Appendix S2**
Figures S3-S6

**Appendix S3**

**Appendix S4**
Tables S2-S5
Figure S7-S8

**Appendix S5**
Figure S9

**Appendix S6**
Figure S10

**Appendix S7**
Figure S11

**Supplementary Tables**
Table S6

**Supplementary Figure**
Figure S12

### Appendix S1: Variation in environmental effects throughout the experiment

##### Motivation

The reciprocal transplant experiment of each population was performed at different dates due to the seasonality of each tested host fruit. To test for potential temporal variability in experimental conditions, we used an inbred *D. suzukii* line (WT3; Chiu *et al.*, 2013) as a control. For each date of experiment, we used this inbred line to measure the three traits of interest: oviposition preference, fecundity and offspring performance. Fecundity and offspring performance were measured in the four test media used in our experiments on wild *D. suzukii* populations: three fruit media (blackberry, cherry and strawberry) and a neutral (i.e., fruit-free) medium (so-called “German food” medium; Olazcuaga et al., 2019). Oviposition preference was measured on the same twelve fruit media as in the main study.

##### Methods

For the statistical analyses of these three traits, the same data transformation was applied as in the main text: log-transformed number of eggs laid for both oviposition preference (i.e., choice assay) and fecundity (i.e., no choice assay), and arc-sinus-transformed egg-to-adult survival for offspring performance.

To estimate the importance of the temporal variability in experimental conditions throughout the different dates of the experiments, we compared for each trait the variance across the different dates (hereafter inter-date variance) and the variance within each date (hereafter intra-date variance). Using a linear model including only a fixed effect to control the quality of the different test-media (hereafter test-fruit), the inter-date variance was estimated by a random effect of the date of the experiment and the intra-date variance was estimated by the variance residuals.

We fitted the following linear mixed model on each of the traits of interest $y_{ijk}$:

$$y_{ijk} = test\_fruit_i + date_j + \varepsilon_{ijk} \quad ,$$

where fixed effects included the effect of the *i*th test fruit medium, $test\_fruit_i$, with $i$=1,...,4, for the four media - blackberry, cherry, strawberry and German food - or $i$=1,...,12 for the twelve fruit media used for the oviposition preference experiment. Random effects included the date of the assay, $date_j$, and a random error, $\varepsilon_{ijk}$, assumed to be normally distributed with a mean of zero and variance $\sigma^2_{inter\text{-}date}$ and $\sigma^2_{intra\text{-}date}$, respectively. To account for potential density-dependence effects on egg-to-adult survival, we added the log-transformed initial egg density as a fixed effect ($eggs_l$). For oviposition preference, we added a random effect of the *l*th arena ($arena_l$) to account for the non-independence between egg counts from the same arena. We assumed $arena_l$ to be normally distributed with a mean of zero and variance $\sigma^2_{arena}$.

**Results and conclusion**

For the three traits, the estimate of the inter-date variance was never higher than the estimate of the intra-date variance (Table S1). In other words, the environmental conditions did not vary more over time, than they varied among replicates (Fig. S1, Fig. S2). These results based on an inbred line validate the comparison of the measurements of traits processed on wild flies collected from different fruits at different dates.

**Table S1. Variance analysis for the three traits of interest i.e., oviposition preference, fecundity and offspring performance.** Estimates of intra-date and inter-date variances, as well as of the variance across the arena for oviposition preference, are shown.

| | **Oviposition preference** | **Fecundity** | **Offspring performance** |
|---|---|---|---|
| $\sigma^2_{inter\text{-}date}$ | 0.250 | 0.094 | 0.029 |
| $\sigma^2_{intra\text{-}date}$ | 1.091 | 0.168 | 0.039 |
| $\sigma^2_{arena}$ | 0.317 | - | - |

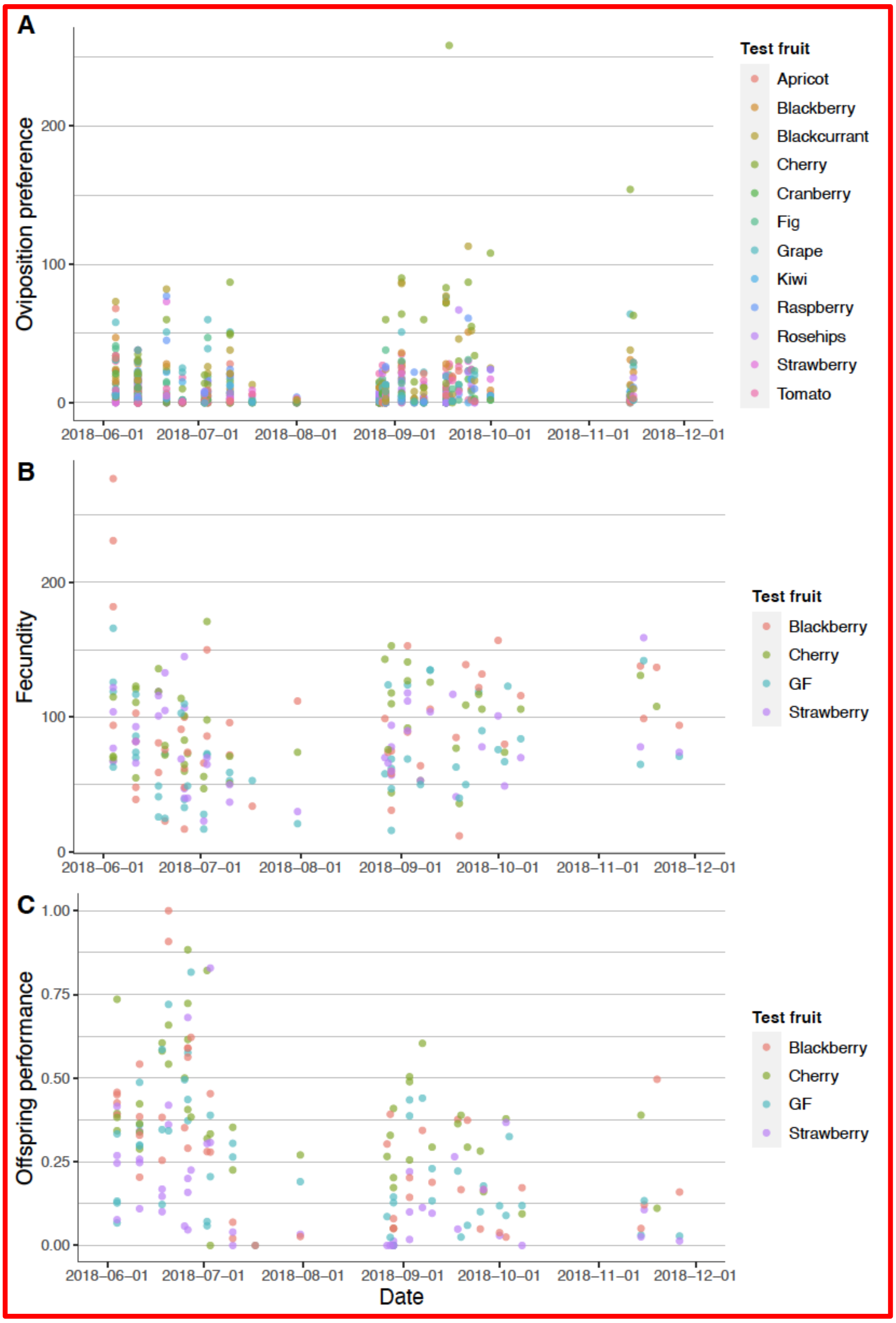


**Figure S1. (A) Oviposition preference (number of eggs laid by 20 females in 24 hours in a choice environment), (B) fecundity (number of eggs laid by 20 females in 24 hours in a no-choice environment), and (C) offspring performance (egg-to-adult survival) measured in the *D. suzukii* inbred line WT3 for each date of reciprocal transplant experiments.**

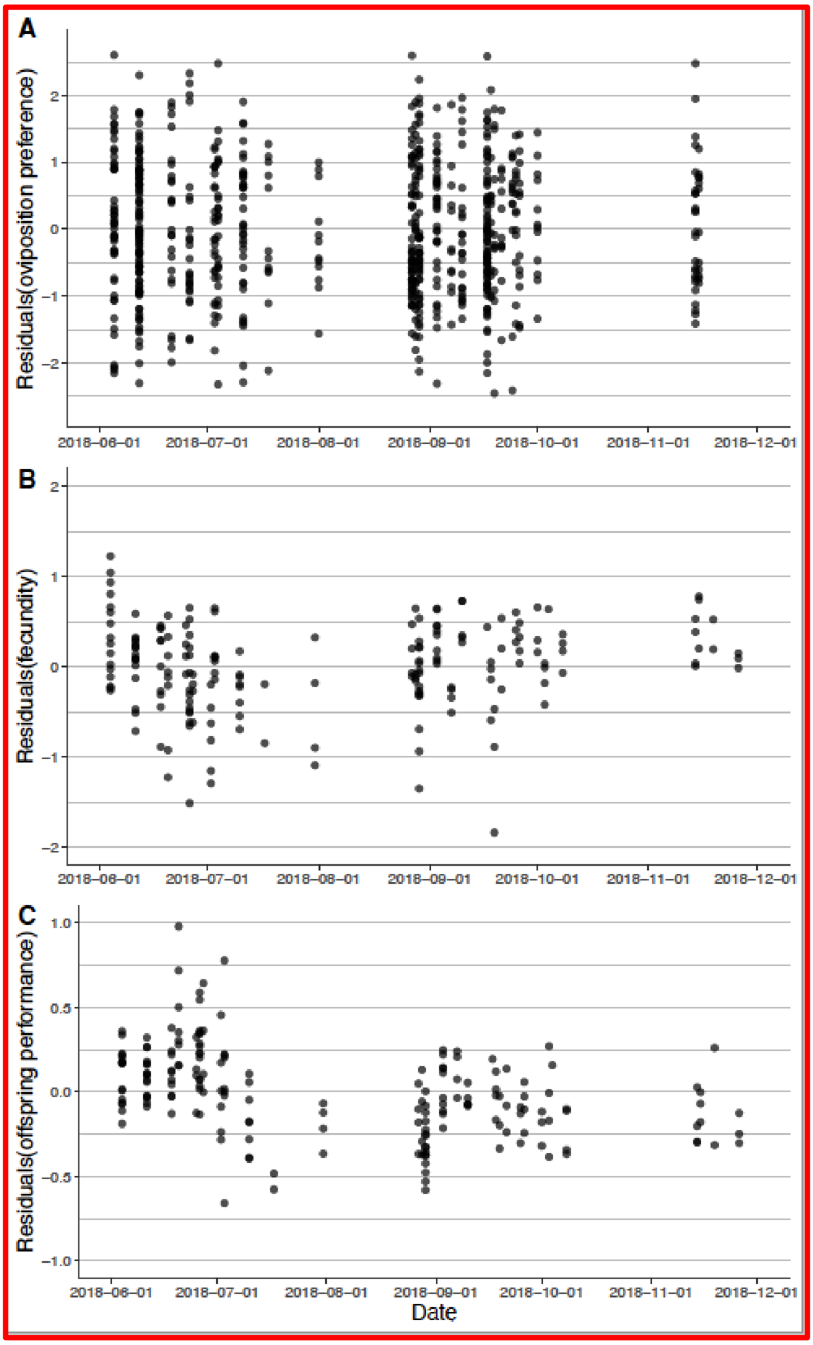


**Figure S2. Residuals of (A) Oviposition preference, (B) fecundity, and (C) offspring performance, measured in the *D. suzukii* inbred line WT3 for each date of reciprocal transplant experiments.** Residual values are estimated after controlling for various confounding factors, including variation in quality among test fruits and among populations for all traits, variation among arenas for oviposition preference or variation among vials with different egg densities for offspring performance.

### Appendix S2: Testing for a pattern of local adaptation by considering the variation in intrinsic quality among test fruits and among populations

#### Motivation and methods

To test whether each trait displayed a pattern of local adaptation we used the method of 'sympatric–allopatric' (SA) contrasts (Blanquart *et al.*, 2013). Briefly, this method tests whether the performance of populations in the original fruit is on average higher than their performance in alternative fruits. Most importantly, the method controls for variation in performance due to variation in intrinsic quality among test fruits and populations.

To implement Blanquart's approach, we fitted the following ANOVA model for each trait of interest (i.e., oviposition preference, fecundity and offspring performance) and this for data obtained in each generation , $y_{ijkl}$ :

$$y_{ijkl} = population_i + test\_fruit_j + SA_{jk} + test\_fruit{\times}original\_fruit_{jk} + \varepsilon_{ijkl} \qquad (2),$$

where fixed effects included the effect of the *i*th population ($population_i$, with i=1,...,25), the effect of the *j*th test fruit ($test\_fruit_j$, with j=1,..., 3, for blackberry, cherry and strawberry, respectively), the interaction between the *j*th test fruit and the *k*th original fruit where the *i*th population was sampled ($original\_fruit_{jk}$, with k=1,..., 3, for blackberry, cherry and strawberry respectively), a sympatric *vs.* allopatric effect that accounts for local adaptation ($SA_{jk}$) and a random error ($\varepsilon_{ijkl}$ with a mean of zero and a variance $\sigma^2_{res}$). We did not include an *original_fruit* effect (which only appears in the interaction described above) as such effect is already statistically accounted for by the population fixed effect. For offspring performance, we accounted for potential density-dependence effects by adding the log-transformed initial egg density as a fixed effect ($eggs_l$). For oviposition preference, we added a fixed effect of the lth arena ($arena_l$) to account for the non-independence between egg counts from the same arena.

To test for a significant pattern of local adaptation, we used a two-way ANOVA and compared the (observed) value of the test-statistics *SA* computed following Blanquart et al. 2013 (eq. D1 in Blanquart et al. 2013's supplementary information) against its expected Fisher-Snedecor distribution with the appropriate degrees of freedom (i.e., F-test). In other words, we tested whether the term $SA_{ijk}$ explained a significant fraction of the variance of the test_fruit×original_fruit interaction (i.e., among each combination of test and original fruits).

Finally, to further visualize variation in quality among populations, we performed a Principal Component Analysis for each trait and each generation, separately.

**Results**

Our analyses show that the power to detect local adaptation or adaptive phenotypic plasticity when using raw data can be reduced by variations in quality among fruit media and among populations (Fig. S3-S5). For example, before the common environment (i.e,. in G0/G1), populations from blackberry consistently had the lowest fecundity and highest offspring performance across the three fruits (Fig. S3C,E). In addition, variation among test fruits is also important. For example, after the common environment (i.e., in G2/G3), all populations had both lower fecundity and lower offspring performance in strawberry medium than in other media (Fig. S3D,F).

In addition, we found that without controlling for variation in intrinsic quality among populations, the increase of a trait value in a fruit is associated with the increase of the trait value in the other fruits for the three traits of interest (Fig. S3).

As emphasized in Blanquart et al. (2013), variations in intrinsic quality among populations and among test fruits can mask patterns of local adaptation and adaptive phenotypic plasticity. This is illustrated when comparing the raw data-based Figure S3 and S4, where variation in intrinsic quality among populations and among test fruits have not been taken into account, with Fig. S5, where such variation has been controlled and residual variation has been computed and represented. The later figure reveals a much clearer pattern of local adaptation in oviposition preference and offspring performance (Fig. S5B,F).

For illustrative purposes, Fig. S6, which is based on a Principal Component Analysis, provides a visual representation of the variation in quality among populations for each trait and each generation.

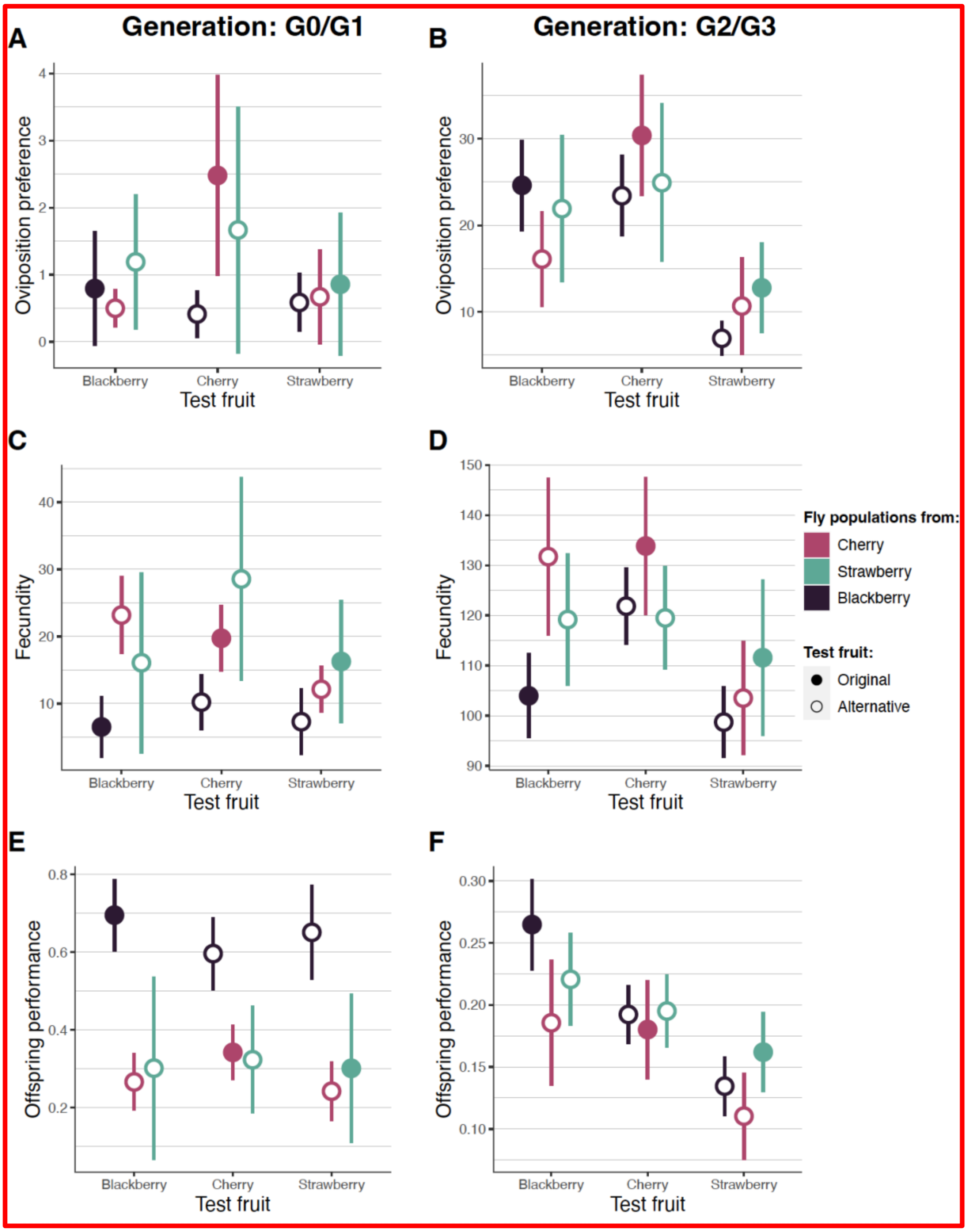


**Figure S3. (A-B) Oviposition preference (number of eggs laid by 20 females in 24 hours on the original or alternative fruit, in a choice environment), (C-D) fecundity (number of eggs laid by 20 females in 24 hours on the original or alternative fruit, in a no-choice environment), and (E-F) offspring performance (egg-to-adult survival) of *D. suzukii* natural populations for each combination of original fruit and test fruit. Here** we used the raw data (without accounting for variation in intrinsic quality among populations and among environments) either before (G0 or G1, depending on the trait measured; left panel) or after the common environment (G2 or G3, right panel). Error bars denote 95% confidence intervals. Sample sizes are 9, 3 and 13 populations assayed for cherry, strawberry and blackberry, respectively.

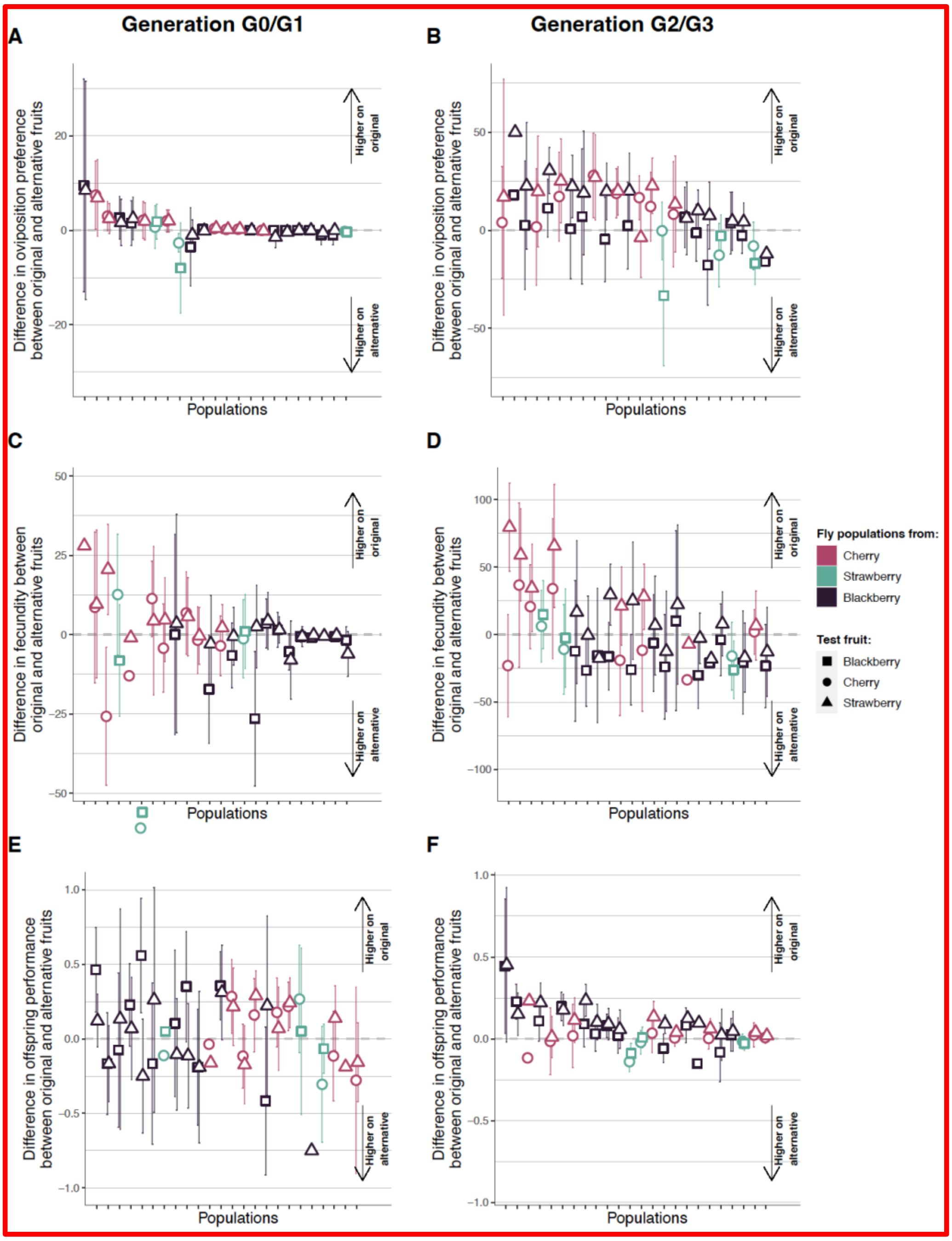


**Figure S4. Difference in (A-B) oviposition preference (number of eggs laid by 20 females in 24 hours in a choice environment), (C-D) fecundity (number of eggs laid by 20 females in 24 hours in a no-choice environment), and (E-F) offspring performance (egg-to-adult survival) between original and alternative fruits for each population in generations G0 or G1 (left panels) and G2 or G3 (right panels).** The difference between original and alternative test fruit for each population was calculated (using raw data) as the average trait value for the original fruit minus the average trait value of alternative fruits. This difference is represented by a symbol whose color depends on the original fruit and shape depends on the test fruit. Populations are ordered following their average trait value measured on the original fruit. Error bars denote 95% confidence intervals.

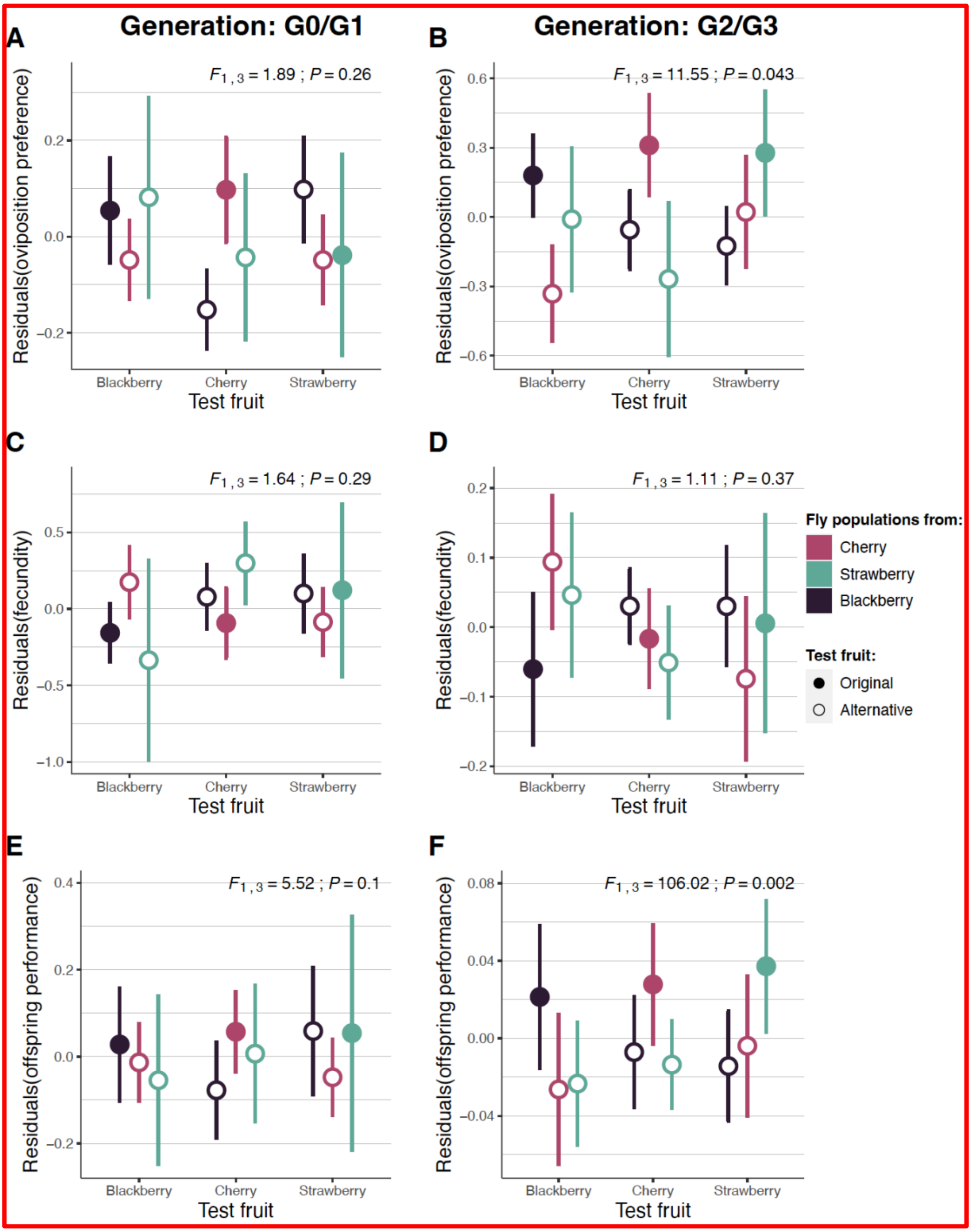


**Figure S5. Residual variation after statistically controlling for intrinsic differences in environment and population quality of (A-B) oviposition preference, (C-D) fecundity, and (E-F) offspring performance of *D. suzukii* natural populations for each combination of original fruit and test fruit either before (G0 and G1, left panel) or after the common environment (G2 and G3, right panel).** For each trait, the mean of residual variations is estimated after controlling confounding factor sources, including variation in quality among test fruits and among populations for all traits, as well as variation among arenas for oviposition preference or among vials with different egg densities for offspring performance. By representing the residuals of a model including a test fruit effect, we control for the quality effect of each test fruit measured across all populations. The representation of these residuals for each population as a function of test fruit thus illustrates the variation of each population according to its original fruit after controlling for these average quality effects. Error bars denote 95% confidence intervals. Sample sizes are 9, 3 and 13 populations assayed for cherry, strawberry and blackberry, respectively.

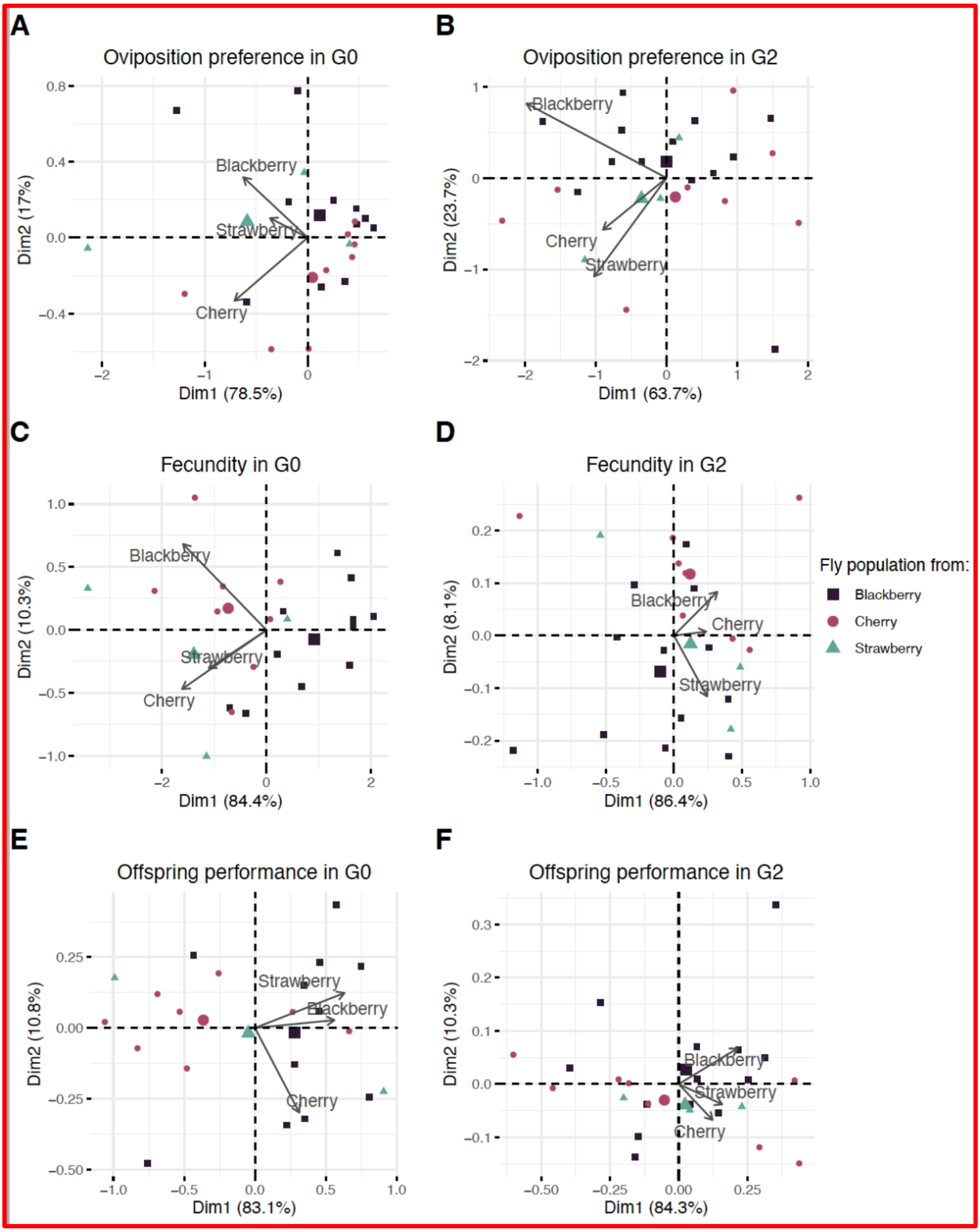


**Figure S6. First and second axes of the Principal Component Analysis (PCA) of all populations across the three test fruits for (A-B) oviposition preference, (C-D) fecundity, and (E-F) offspring performance in generations G0/G1 (left panels) and G2/G3 (right panels).** The mean of each population in each test fruit is represented by a symbol, whose color represents the fruit from which the population originated. The mean is estimated after controlling for variation in quality among fruits (as well as variation among arenas for oviposition preference or variation in the egg density among vials for offspring performance).

### Appendix S3: Estimation of genetic and plastic effects for each combination of test fruit x original fruit

To graphically represent genetic effects and plastic effects for each trait and for each combination of test fruit and original fruit, we used a two-step procedure. First, for each generation separately, we computed the quantity $\varepsilon_{ijk}$ which corresponds to the residuals from a model that only included population and test fruit effects, as well as an arena effect for oviposition preference or an egg number effect for offspring performance. Second, for each combination of test fruit *j* and original fruit *k*, we computed $\underline{\varepsilon_{G0_{jk}}}$and $\underline{\varepsilon_{G2_{jk}}}$ as the average of the residuals in G0 and G2, respectively. Let $N_{G0_{jk}}$ and $N_{G2_{jk}}$represent the number of observations for each combination of test fruit and original fruit in G0 and G2, respectively. The mean of genetic effects corresponds to $\underline{\varepsilon_{G2_{jk}}}$ and its 95% confidence interval can be computed as ± 1.96 x $\frac{\sqrt{\sigma^2_{\varepsilon_{G2_{jk}}}}}{\sqrt{N_{G2_{jk}}}}$. The mean of plastic effects can be computed as the difference between the average of G0 and G2 residuals, $\underline{\varepsilon_{G0_{jk}}} - \underline{\varepsilon_{G2_{jk}}}$, and its 95% confidence interval can be computed as ± 1.96 x$\frac{\sqrt{\sigma^2_{\varepsilon_{G0_{jk}}} - \sigma^2_{\varepsilon_{G2_{jk}}}}}{\sqrt{N_{G0_{jk}}}}$.

### Appendix S4: Statistical power of a new test to detect local adaptation and adaptive phenotypic plasticity based on the $F_{genetic}$ and $F_{plastic}$ tests

#### Motivation

Reliable statistical methods to estimate and test for the relative importance of genetic and plastic effects when a trait has higher value in sympatry than in allopatry are lacking. In our study, we developed a new statistical test to estimate and test for genetic and plastic effects. Our statistical test is based on a modified version of the *F*-test proposed by Blanquart *et al.*, (2013). In the following, we evaluated the statistical power, including false positive rates, of this new test.

#### Methods

- **Simulations**

Data over multiple-generation placed in different test environments after a common environment are required to discriminate genetic from plastic fitness effects. We assume that *N* is the number of original environments (e.g., different types of fruits) in each of which *n* populations are sampled. We simulated fitness data from reciprocal common environment experiments carried out with *m* individuals sampled at two different generations (i.e., before and after a common environment) and transplanted for each combination of test and original environments. In the first generation (G0), we simulated fitness with genetic effects, plastic effects and random environmental effects. In the third generation (G2), we simulated fitness with only genetic effects and random environmental effects.

In G0, the fitness, $y_{ijkl}$, of individuals from the population *i* sampled in the original fruit *k* and placed in the test fruit *j* is simulated as follows:

$$y_{ijkl} = population_{G0G2\_i} + test\_fruit{\times}original\_fruit_{G0\_jk} + test\_fruit{\times}original\_fruit_{G0G2\_jk} + \varepsilon_{ijkl} \quad (3),$$

where the interaction term $test\_fruit{\times}original\_fruit_{G0G2\_jk}$ (respectively $test\_fruit{\times}original\_fruit_{G0\_jk}$) corresponds to a random genetic effect (respectively a random plastic effect). Those both dimensions are equals to $N$, the number of original environments, and whose structures are compound symmetric with variances $\sigma^2_{genetic}$ and covariances $\rho_{genetic}\sigma^2_{genetic}$ (respectively $\sigma^2_{plastic}$ and $\rho_{plastic}\sigma^2_{plastic}$). The $population_{G0G2\_j}$ term is a random population effect assumed to be normally distributed with a mean of zero and variance $\sigma^2_{population}$ and the $\varepsilon_{ijkl}$ term is a random error corresponding to the variation among vials with individuals from the same combination of population and test environment. $\varepsilon_{ijkl}$ is assumed to be normally distributed with a mean of zero and variance $\sigma^2_{res}$.

To simulate both genetic and plastic effects on the difference between sympatric and allopatric combinations, the variances among genetic and among plastic effects ($\sigma^2_{genetic}$ and $\sigma^2_{plastic}$, respectively), the population variance ($\sigma^2_{population}$) and the environmental variance ($\sigma^2_{res}$) were set to values of the same order of magnitude as the ones estimated in our experiment (Table S2). To vary the relative importance of genetic and plastic effects in driving the differences between sympatric and allopatric combinations, the covariances of genetic and plastic effects across environments (i.e., $\rho_{genetic}$ and $\rho_{plastic}$) were set to either -0.5, -0.05 or 0. To simulate the number of eggs in a choice environment, the variance among arenas ($\sigma^2_{arena}$) was set to either 0.5 or 1.5.

In G2, the fitness, $y_{ijkl}$, of an individual from the population $i$ sampled in the original fruit $k$ and placed in the test fruit $j$ is simulated as follows:

$$y_{ijkl} = population_{G0G2\_i} + test\_fruit{\times}original\_fruit_{G0G2\_jk} + \varepsilon_{ijkl} \quad (4),$$

where all terms are as defined above.

To determine whether these statistical tests are robust to different distributions of data, we also simulated data, which follow a Poisson distribution (corresponding to the number of eggs in our study) and a binomial distribution (corresponding to egg-to-adult survival in our study).

First, in G0, the number of eggs, $y_{ijkl}$, laid by individuals from the population *i* sampled in the original environment *k* and placed in the test environment *j* is simulated as follows:

$$y_{ijk} \sim \text{Poisson}\,(\lambda_{ijk}) \qquad (5),$$

where $\lambda_{ijk}$ = exp(population$_{G0G2_i}$ + test_fruit×original_fruit$_{G0_jk}$+ test_fruit×original_fruit$_{G0G2_jk}$) and where all terms are as defined above.

In G2, the number of eggs, $y_{ijkl}$, laid by individuals from the population *i* sampled in the original fruit *k* and placed in the test fruit *j* is simulated as follows:

$$y_{ijk} \sim \text{Poisson}\,(\lambda_{ijk}) \qquad (6),$$

where $\lambda_{ijk}$ = exp(*population*$_{G0G2\_i}$ + *test_fruit×original_fruit*$_{G0G2\_jk}$) and where all terms are as defined above.

In addition, to model the specific context of a choice arena (i.e., number of eggs laid in each square in our study), we also added a random effect of the arena *l* (G0: $\lambda_{ijkl}$ = exp(*population*$_{G0G2\_i}$ + *test_fruit×original_fruit*$_{G0\_jk}$+ *test_fruit×original_fruit*$_{G0G2\_jk}$+ *arena*$_{ijkl}$); G2: $\lambda_{ijkl}$ = exp(*population*$_{G0G2\_i}$ + *test_fruit×original_fruit*$_{G0G2\_jk}$+ *arena*$_{ijkl}$)). The *arena*$_{ijkl}$ term is a random effect, assumed to be normally distributed with a mean of zero and variance $\sigma^2_{arena}$.

Second, in G1, the egg-to-adult survival, $y_{ijk}$, of individuals from the population *i* sampled in the original fruit *k* and placed in the test fruit *j* is simulated as follows:

$$y_{ijk} \sim \text{Binomial}\,(n, p_{ijk}) \qquad (7),$$

where *n* , the initial number of eggs laid in each vial, was assumed to be 20, log($p_{ijk}$ /(1-$p_{ijk}$)) = *population*$_{G0G2\_i}$ + *test_fruit×original_fruit*$_{G0\_jk}$+ test_fruit×original_fruit$_{G0G2_jk}$, and where all terms are as defined above.

In G3, the egg-to-adult survival, $y_{ijk}$, of individuals from the population $i$ sampled in the original fruit $k$ and placed in the test fruit $j$ is simulated as follows:

$$y_{ijk} \sim \text{Binomial}\,(n, p_{ijk}) \qquad (8),$$

where $n$ was assumed to be 20,

$\log(p_{ijk}/(1-p_{ijk})) = population_{G0G2\_i} + test\_fruit{\times}original\_fruit_{G0G2\_jk}$, and where all terms are as defined above.

To evaluate the influence of an unbalanced experimental setup on the statistical power of the test, we first build the design matrix corresponding either to a balanced or an unbalanced experimental design. For the former, the design matrix included 504 observations that correspond to three test environments, three original environments and eight populations sampled in each original environment, and seven individuals assayed for each combination of population and test environment ($N$=3, $n$=8 and $m$=7).

To simulate the design matrix of an unbalanced experimental design, we randomly sampled each row with a probability $p$ from a design matrix corresponding to the balanced design. This process is equivalent to drawing a number between zero and 504 to sample the number of individuals in each combination of population and test environment. The variable $p$ determines the level of unbalance among the number of individuals in each combination and was assumed to follow a dirichlet distribution whose $\boldsymbol{\alpha}$ term was inversely proportional to the level of unbalance in the design. We tested two values of $\boldsymbol{\alpha}$: 1 and 0.25, corresponding to slightly unbalanced and highly unbalanced designs, respectively. A balanced design corresponds to a value of $\boldsymbol{\alpha}$ close to infinity.

**Table S2. Parameters values for the different distributions**

| Distribution | Normal | Poisson | Binomial |
|---|---|---|---|
| $\sigma^2_{genetic}$ | 0.5 | 0.81 | 0.17 |
| $\sigma^2_{plastic}$ | 0.5 | 0.81 | 0.17 |
| $\sigma^2_{population}$ | 0.5 | 0.51 | 0.12 |
| $\sigma^2_{res}$ | 0.5 | - | - |

- **Estimations of $SAplastic_{G0}$ and $SAgenetic_{G0G2}$**

For each of the equations above, we estimated $SAgenetic_{G0G2}$ as the difference between the mean of sympatric values (i.e., average of $test\_fruit{\times}original\_fruit_{G0G2\_jk}$ with $j=k$) and the mean of allopatric values (i.e., average of $test\_fruit{\times}original\_fruit_{G0G2\_jk}$ with $j{\neq}k$). We estimated $SAplastic_{G0}$ similarly, using the values of $test\_fruit{\times}original\_fruit_{G0\_jk}$.

- **Statistical tests**

We tested for genetic and plastic effects using the new SA contrast method described in the main text (Eq 1). As a comparison, we also tested these analyses with linear mixed effects models.

- **Intensity of $SAplastic_{G0}$ and $SAgenetic_{G0G2}$ and statistical power of the tests**

The range of values for $SAgenetic_{G0G2}$ and $SAplastic_{G0}$ varied across statistical distributions (see Results section below). For each distribution, we have kept a large range of $SAgenetic_{G0G2}$ and $SAplastic_{G0}$ values around the absolute values observed in our experiment. We present only the positive values, reflecting an adaptation. Similar patterns were observed with negative SA values reflecting maladaptive processes (not represented here). To estimate the power of the test based on $F_{genetic}$, we kept 500 datasets for each value of $SAgenetic_{G0G2}$

(whatever the value of $SAplastic_{G0}$). We used the same approach for the test based on $F_{plastic}$ and kept 500 datasets for each value of $SAplastic_{G0}$, whatever the value of $SAgenetic_{G0G2}$.

- **False positive rate**

For each of the three data distributions, we estimated the false positive rate of our test, when both $SAgenetic_{G0G2}$ or $SAplastic_{G0}$ are absent, using the same parameters except for the genetic and nongenetic covariances which were set to zero (i.e., $\rho_{genetic} = \rho_{plastic} = 0$). To estimate the false positive rate of the test in the absence of $SAgenetic_{G0G2}$, but when $SAplastic_{G0}$ is present, we used the same parameters except for the genetic covariance which was set to zero (i.e., $\rho_{genetic} = 0$). To estimate the false positive rate of the test in the absence of $SAplastic_{G0}$, but when $SAgenetic_{G0G2}$ is present, we used the same parameters except for the plastic covariance which was set to zero (i.e., $\rho_{plastic} = 0$). We simulated 10,000 datasets for each of these three scenarios.

**Results**

- **Power**

With our simulation parameters, the values of the $SAgenetic_{G0G2}$ and $SAplastic_{G0}$ ranged from 0 to 1 for Normal data, from 0 to 1.5 for Poisson data, and from 0 to 0.4 for Binomial data (Fig. S7-S8). We found that, for both genetic and plastic effects, the statistical power of our test increased with the intensity of local adaptation and adaptive phenotypic plasticity (i.e., with the proportion of variance explained by $SAgenetic_{G0G2}$ and $SAplastic_{G0}$, respectively; Fig. S7). Hence, we found that the $F_{genetic}$ and $F_{plastic}$ tests performed well in a balanced experimental setup.

In the context of an unbalanced experimental setup, we found also that, for both genetic and plastic effects, statistical power increased with the intensity of $SAgenetic_{G0G2}$ and $SAplastic_{G0}$ (Fig. S7). Under an intermediate level of unbalance, $F_{plastic}$ and $F_{genetic}$ performed in the same way as under a balanced experimental setup. Only very strongly unbalanced experimental setups showed a substantial decrease in power.

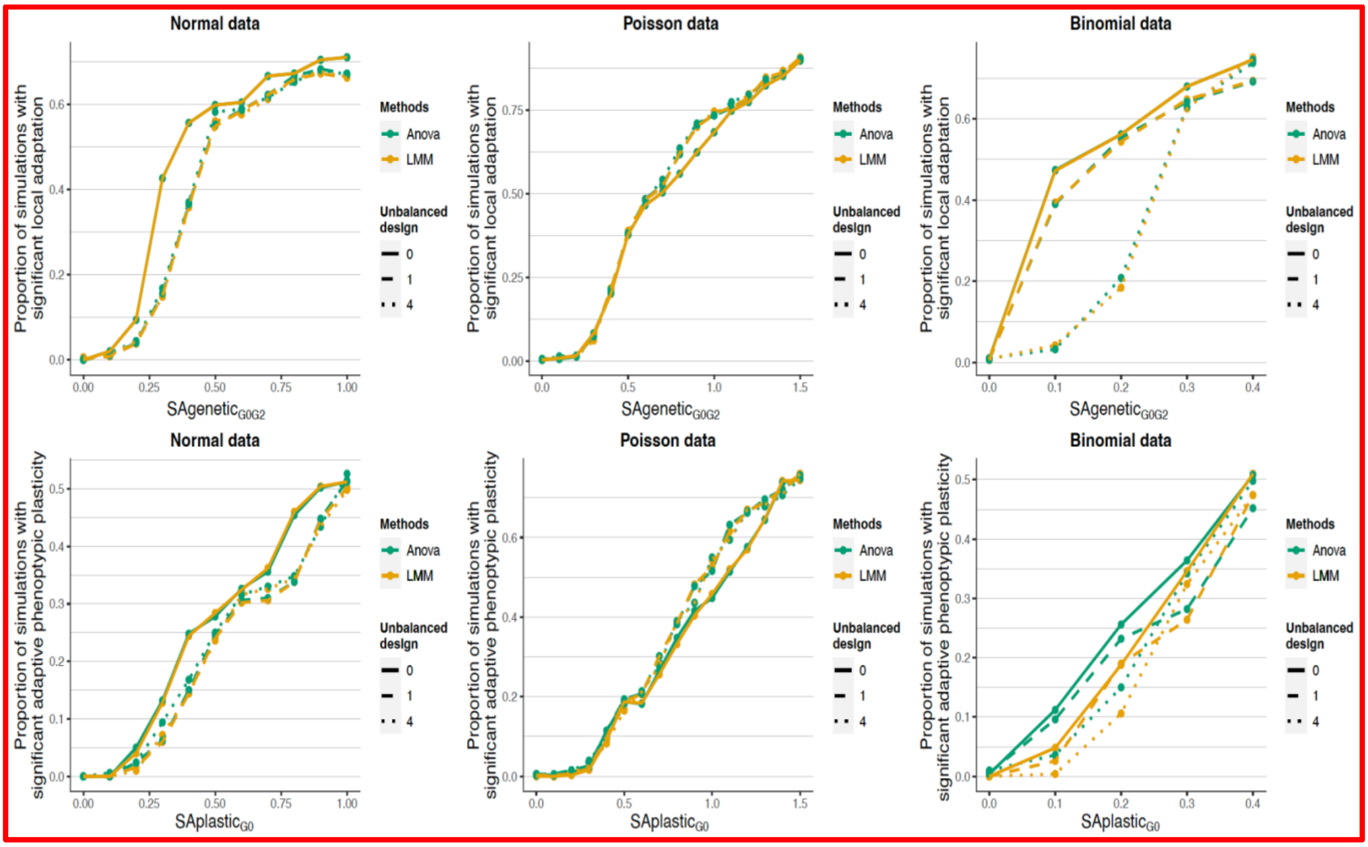


**Figure S7. Power analyses for the tests on both genetic and plastic effects for balanced and unbalanced designs for each of three data distributions: Normal, Poisson and Binomial distribution of data. Anova: *F*-test based on an ANOVA model. LMM: *F*-test based on a Linear Mixed Model (LMM).** For clarity, the level of unbalance of each design is provided as $1/\boldsymbol{\alpha}$.

For oviposition preference, we also verified whether power decreases with increasing variations among arenas. Hence, when we considered a Poisson data distribution with additional variation across arenas, the statistical power for both genetic and plastic SAs decreased as the variance of this arena effect increased (Fig. S8). Nevertheless, $F_{plastic}$ and $F_{genetic}$ tests still performed well even in this case.

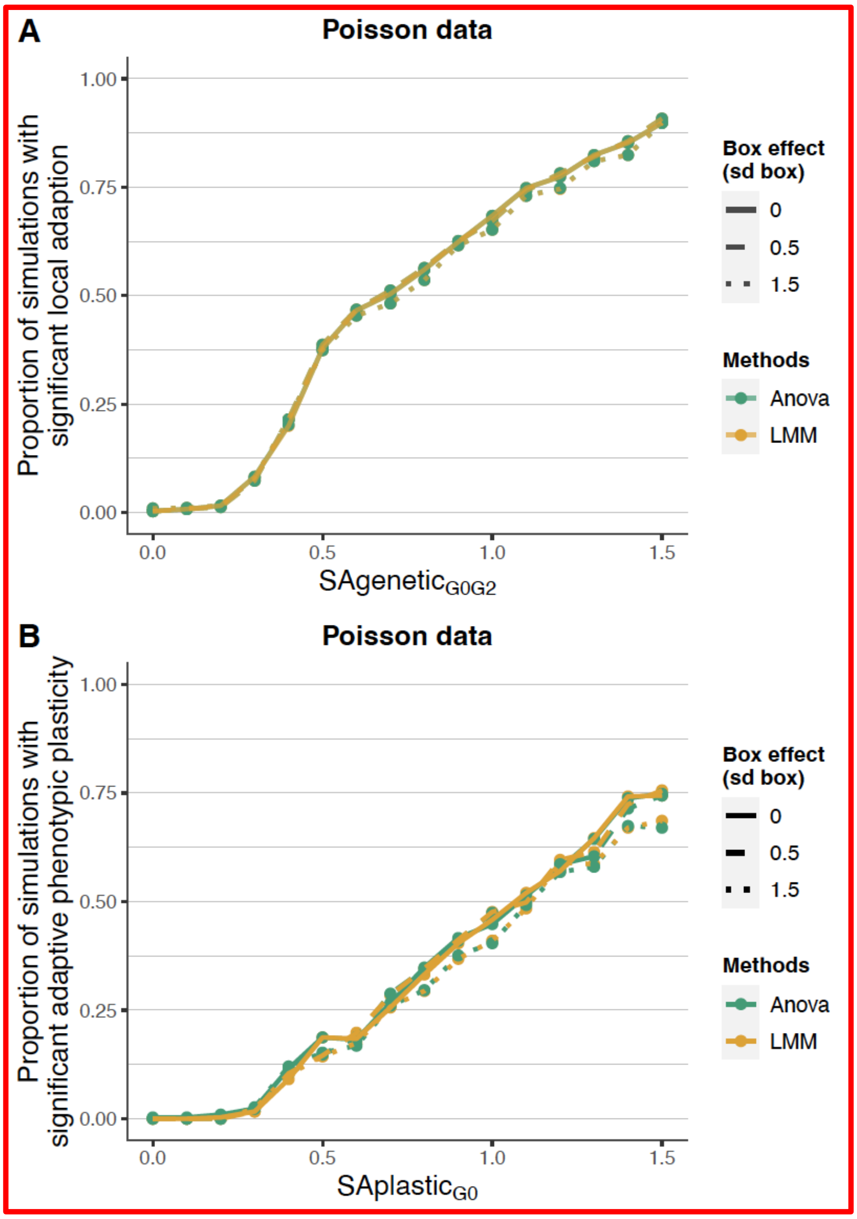


**Figure S8. Power analyses for the tests on both genetic and plastic effects with an arena random effect (Poisson distribution of data)**

- **False positive rate**

With normal data and when either plastic effects, genetic effects or both genetic and plastic effects were absent, the false positive rates ranged from 2.6% to 8.5% (Tables S3, S4 and S5). With Poisson data and binomial data, the false positive rates ranged from 3.0% to 10.8% and from 2.5% to 8.5%, respectively (Tables S3, S4 and S5). While the false positive rates for the detection of $SAgenetic_{G0G2}$ were low in the absence of both $SAgenetic_{G0G2}$ and $SAplastic_{G0}$ (2.9%, 3.2% and 2.5% for Normal, Poisson and Binomial data respectively; Table S3), they were higher when $SAplastic_{G0}$ was greater than zero (8.5%, 10.8% and 8.5%) for Normal, Poisson and Binomial data, respectively (Table S4).

**Table S3. False positive rates for the tests of $SAgenetic_{G0G2}$ and $SAplastic_{G0}$ for each of the three distributions of data.** The absence of both genetic and plastic effects was simulated with $\rho_{genetic}$= 0 and $\rho_{plastic}$= 0.

| Statistical distribution | Normal | Poisson | Binomial |
|---|---|---|---|
| FPR for genetic effects | 2.9% | 3.2% | 2.5% |
| FPR for plastic effects | 2.6% | 3.0% | 2.6% |

**Table S4. False positive rates for the test of $SAgenetic_{G0G2}$ for each of the three distributions of data.** The absence of local adaptation and the presence of adaptive phenotypic plasticity were simulated with $\rho_{genetic}$= 0 and $\rho_{plastic}$= -0.1, respectively.

| Statistical distribution | Normal | Poisson | Binomial |
|---|---|---|---|
| FPR for genetic effects | 8.5% | 10.8% | 8.5% |

**Table S5. False positive rates for the test of $SAplastic_{G0}$ with each of the three distributions of data.** The presence of local adaptation and the absence of adaptive phenotypic plasticity were simulated with $\rho_{genetic}$= -0.1 and $\rho_{plastic}$= 0, respectively.

| Statistical distribution | Normal | Poisson | Binomial |
|---|---|---|---|
| FPR for plastic effects | 2.6% | 4.2% | 2.6% |

**Conclusion**

Our power analysis shows that both $F_{genetic}$ and $F_{plastic}$ tests can be reliably used to test for the relative importance of local adaptation and adaptive phenotypic plasticity, under balanced and unbalanced experimental designs, and for the three tested data distributions (Normal, Poisson and Binomial). Nevertheless, our analysis highlights a false positive rate of ~10% for the detection of genetic effects, when plastic effects are present, and this for the three tested data distributions. It should be noted, however, that when plastic effects are present, the detection of genetic effects can be easily confirmed by testing for significant local adaptation patterns present in G2, but not in G0.

### Appendix S5: Statistical power to detect adaptive phenotypic plasticity with our experimental setup

**Motivation**

While local adaptation was detected, we did not find any (significant) evidence for adaptive phenotypic plasticity (see main text). Since only G0/G1 data were used to detect plastic effects and since the number of individuals available was lower in G0/G1 than in G2/G3, the amount of data used to detect plastic effects was less than half that of genetic effects. The absence of detection of plastic effects could hence be due to a lack of power in our experimental design. Here, we retrospectively performed simulation-based power analyses to determine the power of our experimental design to detect the presence of adaptive phenotypic plasticity. For sake of concision, we focused on offspring performance for this power analysis.

**Methods**

We estimated the power of detecting plastic effects, given that they exist, using simulation-based analyses mimicking our experimental design (Johnson *et al.*, 2015). We simulated data of offspring performance using the estimates of parameters from the model described in Eq. 1 of the main text. In our simulations, all the parameter values were equal to the mean values estimated from our dataset, except for $SAplastic_{G0}$ for which we allowed values to vary from 0 to 1 by a step of 0.10. To evaluate the effect of the number of data, we simulated datasets with different sizes relatively to our original experimental design: datasets X1, X2, X10 and X100 corresponding to datasets with the same size, 2 times bigger, 10 times bigger and 100 times bigger than our original experimental dataset, respectively. We tested for plastic effects using the statistical analyses described in the Statistical methods section of the main text. We simulated 1000 datasets for each value of $SAplastic_{G0}$ and for each type of dataset X1, X2,

X10 and X100. Our statistical power was estimated as the percentage of simulations where we could detect a significant level of adaptive phenotypic plasticity (F-test p-value $< 0.05$).

**Results and conclusion**

As expected, the statistical power increased with the intensity of adaptive phenotypic plasticity (i.e., $SAplastic_{G0}$; Fig. S9) whatever the size of the dataset. Most importantly, the power of our experimental design was roughly similar to that of larger designs (i.e., 2, 10 or 100 times larger datasets). More specifically, our experimental design allows detecting plastic effects with a power greater than an arbitrary threshold of 80% (Johnson et al. 2015) for $SAplastic_{G0}$ values > 0.46, that is when $SAplastic_{G0}$ explains more than 80.4% of the variance of the interaction between the test and original fruits (Fig. S9). In our real *D. suzukii* dataset, the intensity of $SAplastic_{G0}$ was equal to 0.043, a value corresponding to a power of 0.08, with $SAplastic_{G0}$ explaining only 5.94% of the variance of the interaction between the test and original fruits.

In conclusion, if adaptive phenotypic plasticity does exist in our study, its magnitude is likely substantially lower than that we detected for local adaptation in our study and explains less than 80% of the variance of the interaction between the test and original fruits.

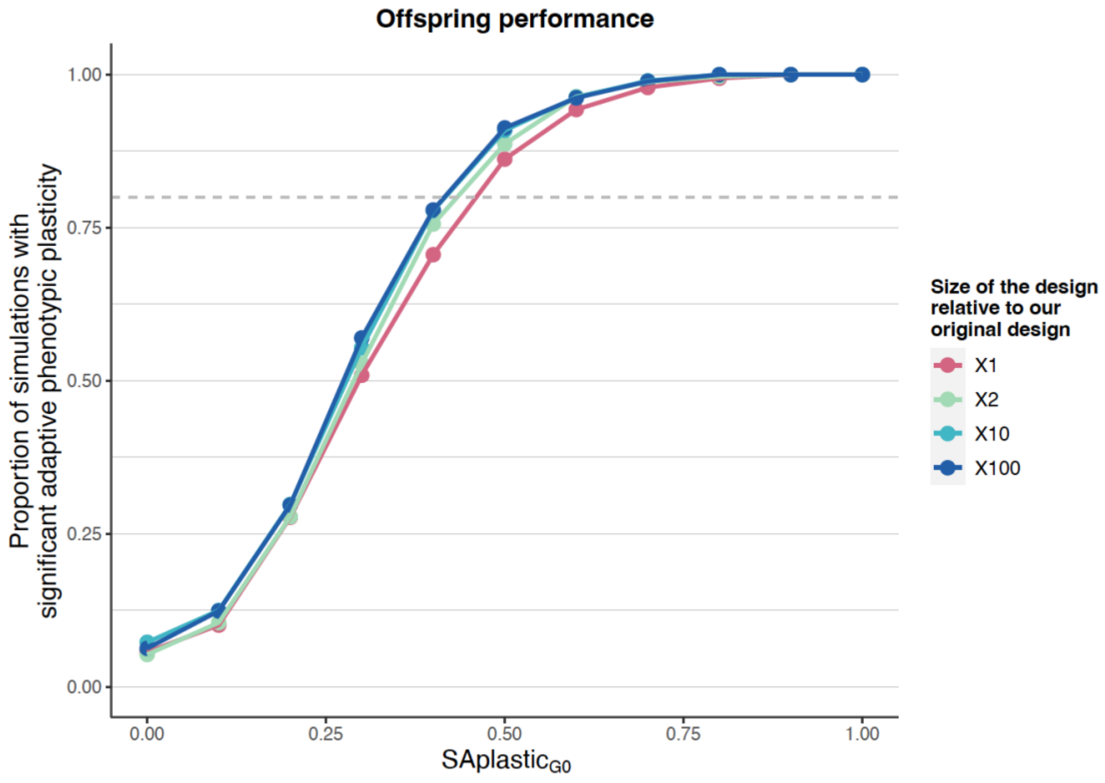


**Figure S9. Estimation of the power of detecting plastic effects for different *SAplastic$_{G0}$* values under our experimental design (X1) and under larger designs (X2, X10 and X100) using simulated data.** Values of *SAplastic$_{G0}$* of 0.05, 0.25, 0.50, 0.75 and 1 explained 6.69%, 45.64%, 79.42%, 90.26% and 94.40% of the variance of the interaction between the test and original fruits, respectively.

### Appendix S6: Correlation across generations for each trait

#### Motivation and methods

To investigate the correlation across generations for each trait, we plotted the average trait value of each population in one generation against its average trait after two generations, after controlling for populations and environment effects as well as other sources of potential other sources of variation, including arena for oviposition preference and egg density for offspring performance.

#### Results and conclusion

Across the three traits of interest and across the three test fruits, the populations with the highest trait value in G0 or G1 did not have the highest trait value in G2 or G3 (Fig. S10). This lack of correlation between generations reflects strong phenotypic plasticity (though not adaptive phenotypic plasticity) in G0 and G1.

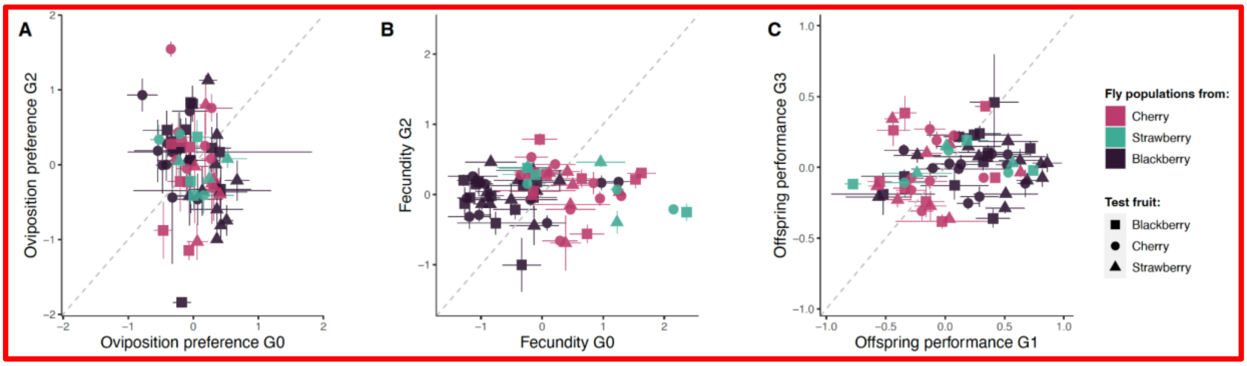


**Figure S10. Relationship between (A) oviposition preference in G0 and G2, (B) fecundity in G0 and G2, and (C) offspring performance in G1 and G3 across the three test fruits.** The mean value of each population in each test fruit is represented by a symbol. The shape and color of the symbols depend on the test fruit and the fruit from which the population originated, respectively. Population mean is estimated after controlling for the quality of each test fruit and the effect of the generation, as well as variation among arenas for oviposition preference or among vials with different egg densities for offspring performance. Error bars represent standard errors of the mean.

### Appendix S7: Test of the "Mother knows best" hypothesis

#### Motivation

The evolution of habitat preference represents an important mechanism for the origin and maintenance of locally adapted phenotypes (Jaenike, 1978; Ravigné *et al.*, 2009). Theory predicts that females may evolve a preference for laying eggs in hosts that maximizes the fitness of their offspring, an evolutionary feature known as the "Mother knows best" principle (Thompson, 1988; Jaenike, 1990). Here, we test for a correlation between female oviposition preference and offspring performance in each of the three studied fruit media, after controlling for populations and environment effects as well as other sources of potential other sources of variation, including arena for oviposition preference and egg density for offspring performance. It is worth noting that fecundity may also be taken as a poor proxy for oviposition preference in the absence of oviposition preference. We found that fecundity was not significantly and negatively correlated with oviposition preference in G0 and G2, respectively. We therefore did not consider fecundity in our analyses, which focused on adult preference.

#### Results

Offspring performance was positively correlated with oviposition preference (Fig. S11). The correlation coefficient was significantly different from zero for the G2/G3 generation test, whereas the 95% confidence intervals of the correlation coefficient included zero for the G0/G1 generation test.

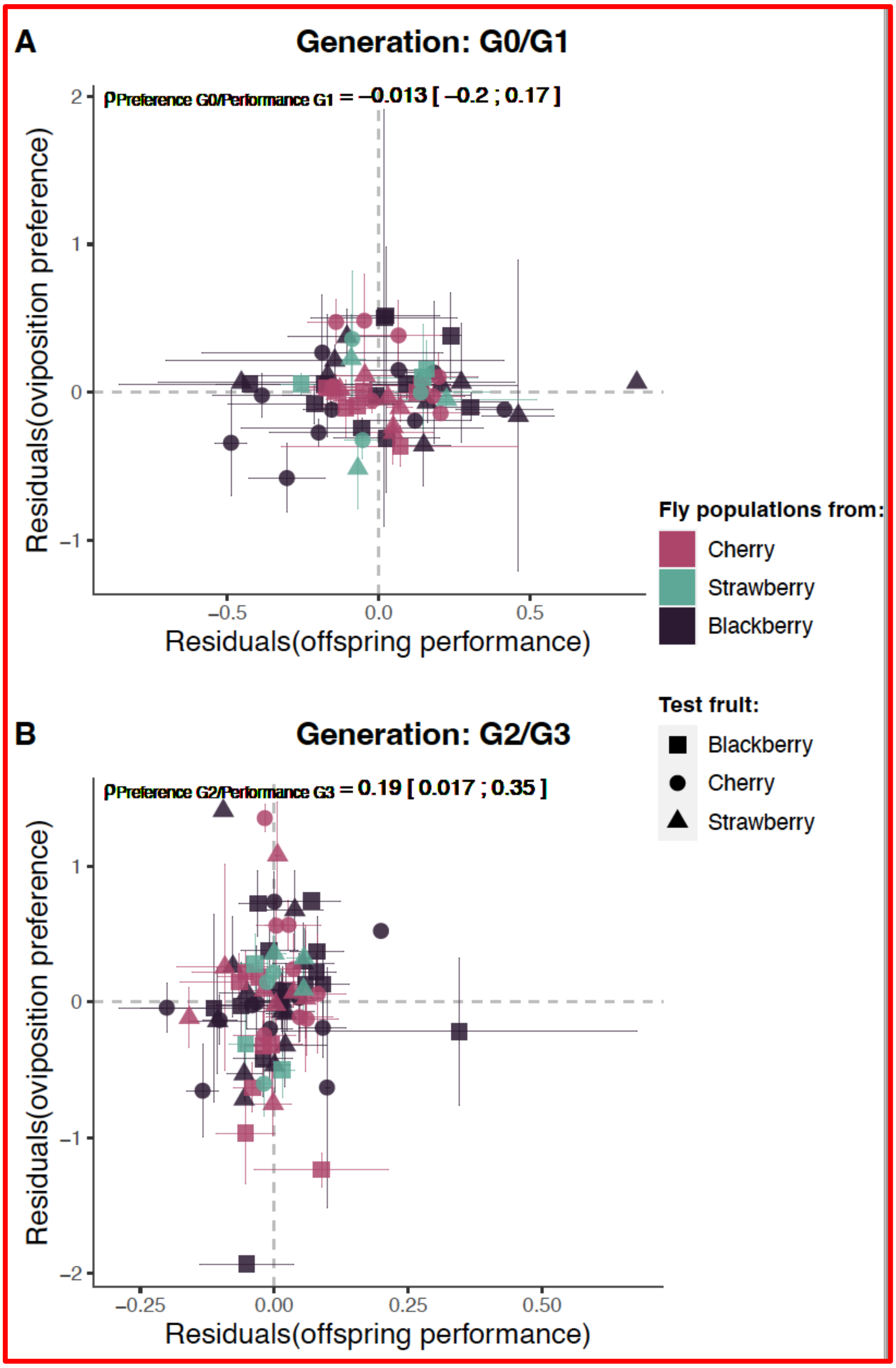


**Figure S11. Correlation between oviposition preference and offspring performance (A) in generation G0/G1 and (B) in generation G2/G3.** The mean trait value of each population in each test fruit is represented by a symbol. The shape and color of the symbols depend on the test fruit and the fruit from which the population originated, respectively. For each trait, the mean of residual variations is estimated after controlling for other sources of variation, including variation in quality among test fruits and among populations for all traits, as well as variation among arenas for oviposition preference or among vials with different egg densities for offspring performance. Error bars represent standard errors of the mean. Correlation coefficients are represented above each panel with the symbol $\rho$ and the associated 95% confidence intervals between brackets.

### Supplementary Tables

**Table S6: Key information about the 25 *D. suzukii* populations collected in the wild that were subsequently analyzed in the laboratory.** The original fruit is the fruit species which was collected in the field and brought into the lab in cages and from which *D. suzukii* flies emerged and were collected for experimental assays. The population code is the code name used in Fig. 2 of the main text. The population name is the code name used for each population during our lab experiment. The city is the name of the closest French city from the sampled site. The GPS coordinates indicate the exact location where fruits were collected. The sampling date using a mm/dd/yyyy format is the sampling date of fruits in the field.

| Original fruit | Population code | Population name | City | GPS coordinates | Sampling date |
|---|---|---|---|---|---|
| Cherry | MTP1 | Cherry50 | Montpellier | 43.626598, 3.863535 | 5/14/2018 |
| Cherry | AVG1 | Cherry51 | Avignon | 43.956177, 4.838228 | 5/23/2018 |
| Cherry | SQP | Cherry52 | Saint-Quentin-la-Poterie | 44.053988, 4.433953 | 5/28/2018 |
| Cherry | CRT1 | Cherry3 | Céret | 42.503621, 2.747655 | 6/1/2018 |
| Cherry | LBL | Cherry6 | Le Boulou | 42.521786, 2.842217 | 6/1/2018 |
| Cherry | CRT2 | Cherry7 | Céret | 42.498589, 2.752859 | 6/1/2018 |
| Cherry | SCR1 | Cherry47 | Saint-Clément-de-Rivière | 43.703934, 3.855531 | 6/6/2018 |
| Cherry | BDX | Cherry103 | Bédarieux | 43.602079, 3.129426 | 6/22/2018 |
| Cherry | TLB | Cherry104 | Taussac-la-Billière | 43.622699, 3.090652 | 6/22/2018 |
| Strawberry | BCR | Strawberry53 | Beaucaire | 43.784667, 4.544464 | 6/8/2018 |
| Strawberry | CND | Strawberry44 | Candillargues | 43.628055, 4.057500 | 6/14/2018 |
| Strawberry | AVG2 | Strawberry42 | Avignon | 43.926049, 4.840473 | 7/12/2018 |
| Blackberry | MGO | Blackberry32 | Mauguio | 43.609988, 3.994697 | 8/28/2018 |

| | | | | | |
|---|---|---|---|---|---|
| Blackberry | SCR2 | Blackberry33 | Saint-Clément-de-Rivière | 43.693692, 3.838453 | 8/29/2018 |
| Blackberry | MTP2 | Blackberry34 | Montpellier | 43.647906, 3.880657 | 8/29/2018 |
| Blackberry | ISS | Blackberry31 | L'Isle-sur-la-Sorgue | 43.914261, 5.079238 | 9/3/2018 |
| Blackberry | CRN | Blackberry35 | Corneilhan | 43.404636, 3.201121 | 9/4/2018 |
| Blackberry | OVL | Blackberry36 | Ouveillan | 43.288458, 2.958087 | 9/4/2018 |
| Blackberry | LDA | Blackberry37 | Laroque-des-Albères | 42.529681, 2.930698 | 9/6/2018 |
| Blackberry | CRT3 | Blackberry38 | Céret | 42.503268, 2.747621 | 9/6/2018 |
| Blackberry | LSR | Blackberry39 | Le Soler | 42.669927, 2.788251 | 9/6/2018 |
| Blackberry | MSL | Blackberry40 | Montferrier-sur-Lez | 43.668816, 3.867852 | 9/8/2018 |
| Blackberry | SGF | Blackberry43 | Saint-Gély-du-Fesc | 43.669139, 3.839824 | 9/11/2018 |
| Blackberry | BKZ | Blackberry45 | Blauzac | 43.953888, 4.381957 | 9/12/2018 |
| Blackberry | LBS | Blackberry44 | La Boissière | 43.667453, 3.660270 | 9/13/2018 |

### Supplementary Figures

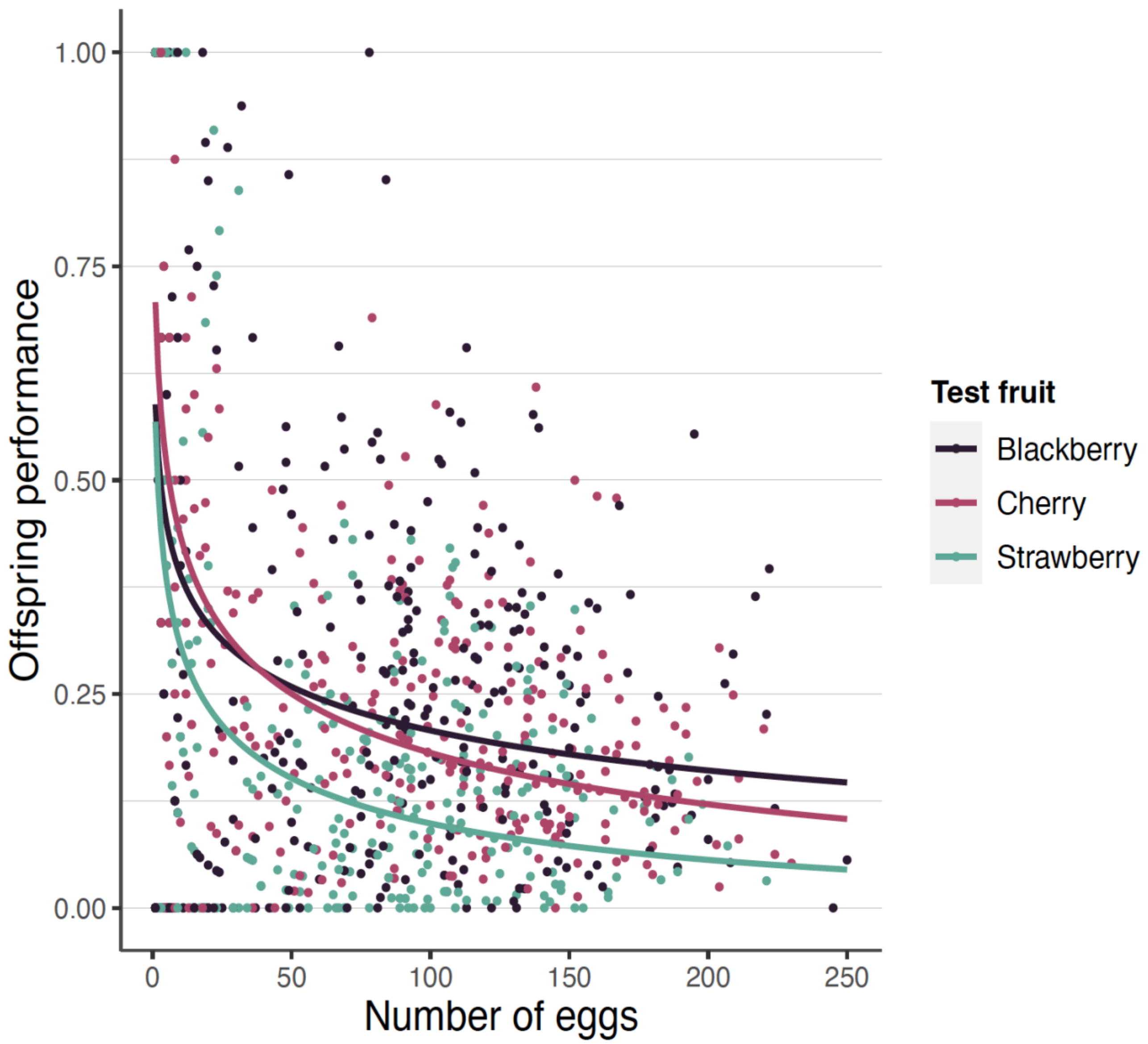


**Figure S12. Effect of egg density on offspring performance for each test fruit.** Observed data are shown as small dots. The linear regression of the arc-sine-transformed egg-to-adult survival explained by the log-transformed egg density for each fruit media is shown as a solid line.
